# A dual enhancer-silencer element ensures transient *Cdx2* expression during posterior body formation

**DOI:** 10.1101/2024.04.10.588878

**Authors:** Irène Amblard, Damir Baranasic, Benjamin Moyon, Boris Lenhard, Vicki Metzis

**Affiliations:** Institute of Clinical Sciences, Faculty of Medicine, Imperial College London, W12 0HS, UK; Medical Research Council (MRC) Laboratory of Medical Sciences (LMS), London W12 0HS, UK; Division of Electronics, Ruder Boskovic Institute, Bijenicka cesta 54, 10000 Zagreb, Croatia

**Keywords:** silencer, enhancer, motif composition, WNT signalling, retinoic acid nuclear receptor, CDX, spinal cord development, directed differentiation of mouse embryonic stem cells

## Abstract

During development, cells express precise gene expression programmes to assemble the trunk of the body plan. Appropriate control over the duration of the transcription factor *Cdx2* is critical to achieve this outcome, yet, how cells control the onset, maintenance or termination of *Cdx2,* has remained unclear. Here, we delineate the cis-regulatory logic orchestrating dynamic *Cdx2* expression in mouse caudal epiblast progenitors and their derivatives - spinal cord and presomitic mesoderm. Combining CRISPR-mediated deletion of regulatory elements with *in vitro* models and *in vivo* validation, we demonstrate that distinct enhancers, and a silencer, embedded at the *Cdx2* locus, act sequentially to drive transient *Cdx2* expression. We pinpoint a minimal silencer element that relies on a nuclear receptor motif to extinguish *Cdx2*. Changing this single motif converts the repressive element to an enhancer with opposite regulatory behaviour. Our findings elucidate design principles of developmental enhancers and silencers and establish a dual enhancer-silencer cis-regulatory logic ensuring precise spatiotemporal control over gene expression for vertebrate body patterning.

## INTRODUCTION

In the mammalian body, a striking array of cell types emerges during development, in response to extrinsic cues. This immense diversity results from the activity of gene regulatory networks that define cell identity^1^. Yet, how cells interpret extrinsic signalling in a context-dependent manner to ensure the generation of different cell types, remains a major open question.

CDX (CDX1, 2 and 4) transcription factors play a central role in the development of the caudal part of the body plan^2–5^. Removal of these partially redundant factors^6–7^ results in the loss of most post-occipital tissues, in part due to their regulation of *Hox* genes^4,8–10^. Reduced or prolonged expression of CDX factors respectively truncates or expands the territory that forms the spinal cord, at the expense of hindbrain fates, in multiple species^6,11–15^. Unlike *Cdx1* and *Cdx4*, however, genetic removal of *Cdx2* alone demonstrates its indispensable role in posterior body formation^3,6–8,16–17^.

The expression of *Cdx2* is detected during gastrulation in the mouse caudal epiblast^18–20^. This region of the embryo harbours neuromesodermal progenitors (NMPs), a progenitor source that contributes to the developing spinal cord and somites^21–23^. Although *Cdx2* is detected in NMPs^24–26^, and is later maintained in the hindgut^18,26^, it is only transiently expressed in tissues derived from NMPs, such as the spinal cord and somites^3,9,26–27^. WNT and FGF signalling promote caudal embryo development, and the expression of *Cdx2*^12–13,28–29^. Similar regulation is observed *in vitro* using the directed differentiation of mouse or human ESCs^30–34^. In addition, retinoic acid (RA) signalling is a critical determinant of posterior body formation and differentiation^35–36^ and restricts the expression of *Cdx2* in the spinal cord *in vivo*^27^ and *in vitro*^24,37^. The transition from a caudal epiblast to a spinal cord progenitor coincides with a switch from FGF to RA signalling^35–36^. How cells coordinate the duration of *Cdx2* expression in response to extrinsic cues, remains unresolved.

Extrinsic cues are interpreted by cis-regulatory elements (CREs) to control gene expression. CREs encompass a broad range of elements that include promoters, enhancers^38^, silencers^39^, and insulators^40^. Recent findings suggest that enhancers may function in concert with addition elements, such as tethers^41^ and more recently, facilitators^42^, yet how common such elements are in the genome remains unclear. Although many classes of CREs exist, the ability to predict the cellular context and function of individual CREs remains challenging^43^. Several *Cdx2* enhancers have been identified that partially recapitulate the trophectoderm, caudal epiblast or intestinal expression pattern of *Cdx2 in vivo*^44–47^. These studies suggest that a subset of *Cdx2* CREs play tissue-specific roles. Although long-range interactions can play a vital role in regulating developmental genes^48–49^, the expression of *Cdx2* during posterior body formation appears to be regulated by elements located within the *Cdx2* locus. In particular, an 11kb region flanking *Cdx2* recapitulates the caudal tailbud expression pattern of *Cdx2* between E7.5 and E10.5^50^. Strikingly, several CREs located within this region demonstrate enhancer activity in transgenic reporter embryos, yet, as individual elements they do not recapitulate the full expression pattern of *Cdx2*^47,50^. How multiple CREs within their native genomic context facilitate *Cdx2* initiation, maintenance or termination, remains unclear.

In this study, we investigate the regulatory mechanisms responsible for controlling the duration of *Cdx2* during the formation of posterior body derivatives: spinal cord (SC) and paraxial presomitic mesoderm (PSM) progenitors. To dissect the molecular mechanisms that control *Cdx2*, without compromising axial elongation or trophectoderm specification^51^, we use genome engineering approaches, combined with the directed differentiation of pluripotent ESCs to model posterior body formation *in vitro*. Using this strategy, we provide evidence that the transient expression of *Cdx2* in cells relies on the sequential usage of CREs that perform discrete, nonredundant functions during development. We identify the location of a *Cdx2* silencer, and demonstrate that the substitution of a single retinoic acid nuclear receptor binding motif can convert this element into an enhancer. Furthermore, we validate that silencer function is mediated by this motif *in vivo*, demonstrating that mutations within the motif impact CDX2 and the resulting regional identity in mouse embryos. Taken together, we provide evidence that the composition and number of retinoic acid nuclear receptor motifs dictates silencer versus enhancer function, and underpins the context-specific regulation of *Cdx2* during posterior body development.

## RESULTS

### *Cdx2* CREs display transient accessibility during spinal cord development

To identify putative CREs regulating *Cdx2* expression during posterior body formation, we examined the chromatin accessibility landscape inferred from ATAC-seq experiments using two different approaches. We complemented an *in vitro* time course of bulk ATAC-seq data from mouse embryonic stem cells (ESCs) differentiated into spinal cord (SC) progenitors, which transiently express *Cdx2* (Fig1A-C)^52^, with pseudo-bulk ATAC-seq profiles obtained from the corresponding cell types present *in vivo*, extracted from 10x multiome single-nucleus (sn)ATAC-seq performed on E7.5-E8.75 whole embryos (Fig1C)^53^. The transient expression of *Cdx2 in vitro* corresponds to caudal epiblast-like (CEpiL) cells, which upon differentiation to SC, lose *Cdx2*^31,52,54^ (Fig1A-B; see methods). As genome-wide changes in chromatin accessibility take place in CEpiL versus SC progenitors^52,54–55^, we hypothesized that changes in the availability of CREs may contribute to the regulation of *Cdx2*.

**Figure 1:**
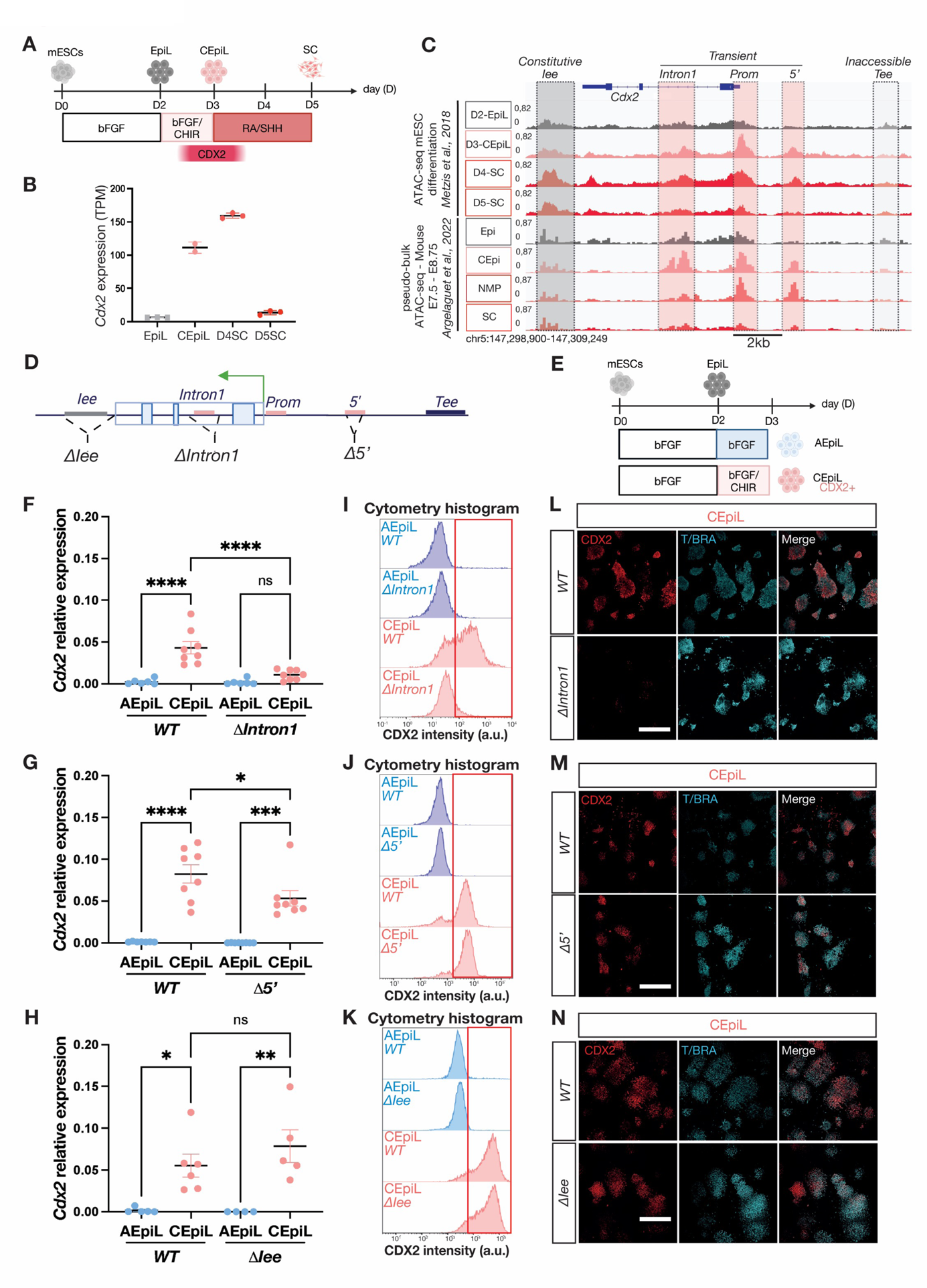
An intronic enhancer is indispensable for the onset of *Cdx2* in caudal epiblast conditions. **(A)** Conditions used to generate spinal cord progenitors from mouse embryonic stem cells over a 5-day period (see 52). (**B**) *Cdx2* expression detected using mRNA-seq during the directed differention of ESCs to spinal cord progenitors. Data reprocessed from Gouti et al., 2014 (31) (see methods). TPM= transcripts per million. (**C**) ATAC-seq signal at the *Cdx2* locus between day 2-5 of the differentiation in A. Single-nucleus pseudobulk ATAC-seq signals at the *Cdx2* locus showing accessibility in epiblast (Epi), caudal epiblast (CEpi), neuromesodermal progenitors (NMP) and spinal cord (SC) from 53. The promoter (*Prom*), *Intron1* (50), and *5’* elements (50) gradually lose accessibility as cells progress to day 5 spinal cord (regions shaded in pink) in contrast to the accessible intestinal enhancer (*Iee*, 47) (grey shading) and inaccessible trophectoderm enhancer (*Tee*; 44), (grey box; no shading) **(D)** Schematic of the *Cdx2* locus illustrating the position of elements targeted for removal in mouse ESCs using pairs of CRISPR (see methods). ESCs harbouring deletions correspond to either the Iee *(ΔIee*); first intron *(ΔIntron1*), 5’ (*Δ5’*) or TEE (*ΔTee*). **(E)** Conditions used to generate caudal epiblast-like (CEpiL) cells *in vitro* that express *Cdx2* in response to a brief pulse of bFGF and CHIR99021 between day 2-3 (pink), versus anterior epiblast-like (AEpiL) cells used as a control that are only exposed to bFGF (blue) and do not express *Cdx2*. **(F-H)** Relative expression (RT-qPCR) for *Cdx2* in AEpiL (blue) versus CEpiL (pink) conditions from collected from *ΔIntron1* (F, n=8), *Δ5’* (G, n=8) and *ΔIee* (H, n=6) cells demonstrates that *Cdx2* is not induced in CEpiL cells that lack the *Intron1* element (G, p<0.0001, Tukey’s test) while the removal of the *5’* element result in a slight decrease (H, p=0.0436; Tukey’s test). (**I-K**) Representative flow cytometry histograms for CDX2 in AEpiL (blue) versus CEpiL (pink) conditions collected from *ΔIntron1* (I), *Δ5’* (J) and *ΔIee* (K) cells demonstrates a severe depletion of CDX2 positive progenitors in *Intron1* lacking CEpiL cells (I). **(L-N)** Immunofluorescence for CDX2 and TBXT/BRACHYURY demonstrates that *ΔIntron1* cells (L) maintain expression of TBXT in CEpiL conditions despite loss of CDX2. Scale bar = 500μm. n=3. Panels A, D, E created with BioRender.com

Consistent with this, known *Cdx2* CREs exhibited distinct patterns of chromatin accessibility^44–47,50,56^. In particular, a set of three CREs were transiently accessible *in vitro*, when comparing CEpiL (Cdx2 positive; Fig1B-C) to SC progenitors (Cdx2 lacking; Fig1B-C). These corresponded to the *Cdx2* promoter, an *Intron1* CRE^50^, and a CRE located upstream of the transcriptional start site of *Cdx2*^50^, termed *5’* (Fig1C, “*transient*”, boxed in pink). Similarly, *in vivo*, these regions are accessible in the caudal epiblast (CEpi; Fig1C), but appear inaccessible in SC progenitors (Fig1C). Both the *Intron1* and *5’* CRE exhibit enhancer activity in posterior tissues, although their onset and specificity differ to *Cdx2*^47,50^. By contrast, CREs that regulate *Cdx2* in other tissues such as the intestine (*Iee*) or trophectoderm (*Tee*), appear either continuously accessible (“*constitutive*”, Fig1C; box shaded in grey) or largely lacking accessibility, respectively, in the same cellular conditions examined *in vitro* and *in vivo* (“*inaccessible*”, Fig1C; white box). In summary, we show that a defined set of *Cdx2* CREs^50,56^ are transiently accessible at the time *Cdx2* is expressed, in an *in vitro* model of SC development.

### Separate CREs control the onset versus maintenance of *Cdx2* expression

Having established that a distinct set of *Cdx2* CREs are transiently accessible, we set out to test directly the function of each CRE on the regulation of *Cdx2* during posterior body formation using a previously established ESC *in vitro* system to model spinal cord or paraxial mesoderm development^24,54^. We engineered a suite of ESC lines that lacked individual CREs corresponding to either *transient*, *constitutive*, or *inaccessible* regions using CRISPR/Cas9 mediated genome editing (Fig1D and FigS1A; and methods). WT versus CRISPR mutant ESC lines were then differentiated towards CEpiL cells that express *Cdx2* in response to a brief pulse of bFGF and CHIR ^24,31,52,57^ (methods). Anterior epiblast-like (AEpiL) cells, which do not express *Cdx2*, were used as a control and induced by exposure to bFGF alone (see methods) (Fig1E and FigS1B)^31,33,52,54^. *Cdx2* induction was assayed by RT-qPCR and immunofluorescence (IF), together with flow cytometry, to investigate CDX2 in a quantitative and single-cell manner. The expression of CDX2 remained indistinguishable between WT cells and cells lacking either the *5’* (Fig1G,J,M and FigS1G), the *Iee* (Fig1H,K,N and FigS1H) or the *Tee* (FigS1C,E,I) CRE *(referred to as Δ*5’, *ΔIee* and *ΔTee* cells, respectively). By contrast, removal of the *Intron1* CRE severely impaired the induction of *Cdx2*, as demonstrated at the transcript (Fig1F) and protein level (Fig1I,L and FigS1J). Impaired induction of CDX2 was also recapitulated by the removal of the promoter for *Cdx2* (FigS1D,F,K). This demonstrates that the intronic CRE plays an indispensable role in the induction of *Cdx2* in CEpiL cells. These data demonstrate that the removal of the *Intron1* CRE is sufficient to block *Cdx2* induction in CEpiL cells, despite the presence of several alternative, and accessible, regulatory regions at the *Cdx2* locus, such as the *Iee* or the *5’* CRE (Fig1C).

Having established that *Intron1* is indispensable for *Cdx2* induction in CEpiL cells, we tested whether *Intron1* was required for *Cdx2* expression in alternative cell types. To assess this, we differentiated ESCs under presomitic mesoderm (PSM) conditions (Fig2A), in which CDX2 is sustained for a longer time period (day 3-5) relative to spinal cord progenitors (day 3-4) (Fig2A-B). Despite an initial loss of *Cdx2* in CEpiL cells lacking *Intron1,* the expression of CDX2 is recovered at Day 4 to levels comparable to WT cells in PSM conditions (Fig2C-D; D4PSM condition). These data demonstrate that *Intron1* plays a cell-type specific role in regulating *Cdx2*, and suggest that an alternative CRE may be responsible for *Cdx2* expression in PSM conditions.

**Figure 2:**
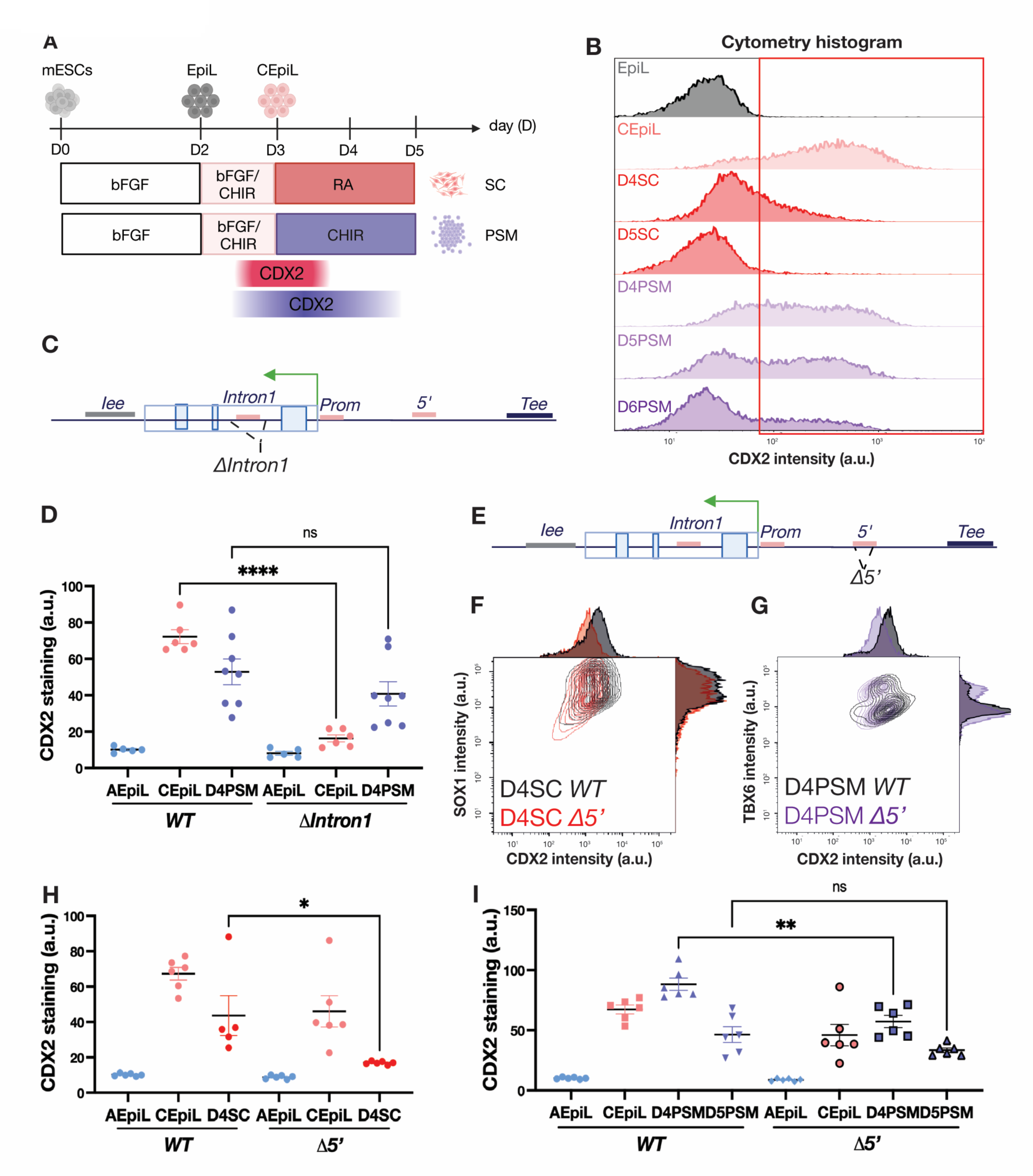
*Cdx2* is transiently maintained in caudal epiblast derivatives via an alternative, 5’ cis-regulatory element. **(A)** Schematic of mouse ESCs differentiated into spinal cord (SC) or presomitic mesoderm (PSM) progenitors highlighting the transient expression of *Cdx2*, which is lost more rapidly in spinal cord (pink) conditions. (**B**) Representative flow cytometry histogram for CDX2 in epiblast (grey), caudal epiblast (CEpiL, pink), day 4 and day 5 SC (red) or day 4, day 5, day 6 PSM (purple) conditions collected from WT cells shows that CDX2 persists longer in PSM conditions. (**C**) Schematic illustrating the *Intron1* element removed in mouse ESCs (*ΔIntron1*). (**D**) CDX2 levels assessed by flow cytometry in anterior epiblast-like (AEpiL, blue), CEpiL cells (pink) and D4 PSM (purple) conditions collected from WT and *ΔIntron1* cells shows comparable CDX2 levels in WT and *ΔIntron1* in D4 PSM cells (n=8; p= 0.2349, two-tailed t-test). (**E**) Schematic illustrating the position of the 5’ element removed in mouse ESCs (*Δ5’*). (**F**) Representative flow cytometry contour and histogram plot for CDX2 (X axis) and SOX1 (Y axis) in D4 SC conditions collected from *WT* (black) and *Δ5*’(red) cells determined by flow cytometry. (**G**) CDX2 levels assessed by flow cytometry in AEpiL (blue), CEpiL (pink) and D4 SC (red) conditions collected from WT and *Δ5’* cells show a significant decrease of CDX2 in D4SC cells (n=5; p=0.0281; two-tailed t-test). (**H**) Representative contour and histogram plot for CDX2 (X axis) and TBX6 (Y axis) in D4PSM conditions collected from *WT* (black) and *Δ5*’ (purple) cells. (**I**) CDX2 levels assessed by flow cytometry in AEpiL (blue), CEpiL (pink) and D4PSM (purple) conditions collected from WT and *Δ5’* cells show a significant decrease in D4 PSM cells (n=6; p=0.0015; two-tailed t-test). Panels A, C, and E created with BioRender.com.

The *5’* CRE demonstrates transient accessibility, and, relative to the *Intron1* CRE, it exhibits a slightly later onset of enhancer reporter activity in the mouse caudal epiblast (50). Removal of the *5’* CRE in ESCs (Fig2E), followed by their differentiation towards SC or PSM (Fig2A) resulted in reduced CDX2 expression in both conditions relative to WT cells (Fig2F-I). RT-qPCR primers designed to detect spliced versus nascent *Cdx2* transcripts confirmed that a reduction in *Cdx2* was detectable at the level of transcription (FigS2D,F). This suggests that *Cdx2* expression in derivatives of the CEpiL cells is dependent on the *5’* CRE, and, upon its removal, *Cdx2* is rapidly downregulated, decreasing the duration of expression (FigS2C-F). Taken together, transient chromatin accessibility changes can be used to predict key regulatory elements that play nonredundant roles in the onset of *Cdx2* expression in CEpiL cells (*Intron1*) versus spinal cord or paraxial mesoderm (*5’*) progenitors.

### The intronic CRE harbours a silencer and an enhancer for *Cdx2*

To investigate what determines CRE usage in different cellular conditions, we sought to define what factors are responsible for CRE activity. CEpiL cells require active WNT signalling conditions to express *Cdx2* ^9,31,33,52^ (Fig1E). We therefore examined the ChIP-seq signal of several WNT effectors in naïve mouse pluripotent ESCs versus CEpiL cells (Fig3A), distinct cellular conditions which respectively repress or promote *Cdx2* expression^54^. Analysis of these data revealed that despite the presence of TCF/LEF sites in multiple, accessible CREs (*Intron1*, *5’*, and *Iee*^47,50^; Fig1C), the occupancy of WNT effectors is highly selective. CTNNB1 (βCAT), LEF1, and TCF3 are exclusively occupying the intronic CRE in both conditions (Fig3A). Further examination of the intronic CRE revealed that WNT effectors occupy two distinct sub-regions within the *Intron1* CRE (labelled *P1* and *P2*, Fig3A), depending on the cellular conditions. In naïve pluripotency conditions, in which *Cdx2* is repressed, *P2* is occupied by CTNNB1 and TCF3; by contrast, the expression of *Cdx2* in CEpiL cells coincides with LEF1 preferentially occupying *P1*, while TCF3, and to a lesser extent, CTNNB1, are depleted at *P2*. We hypothesized from these data that *P1* and *P2* may mediate opposing regulatory functions that favour activation (at *P1*) versus repression (at *P2*) of *Cdx2*.

**Figure 3:**
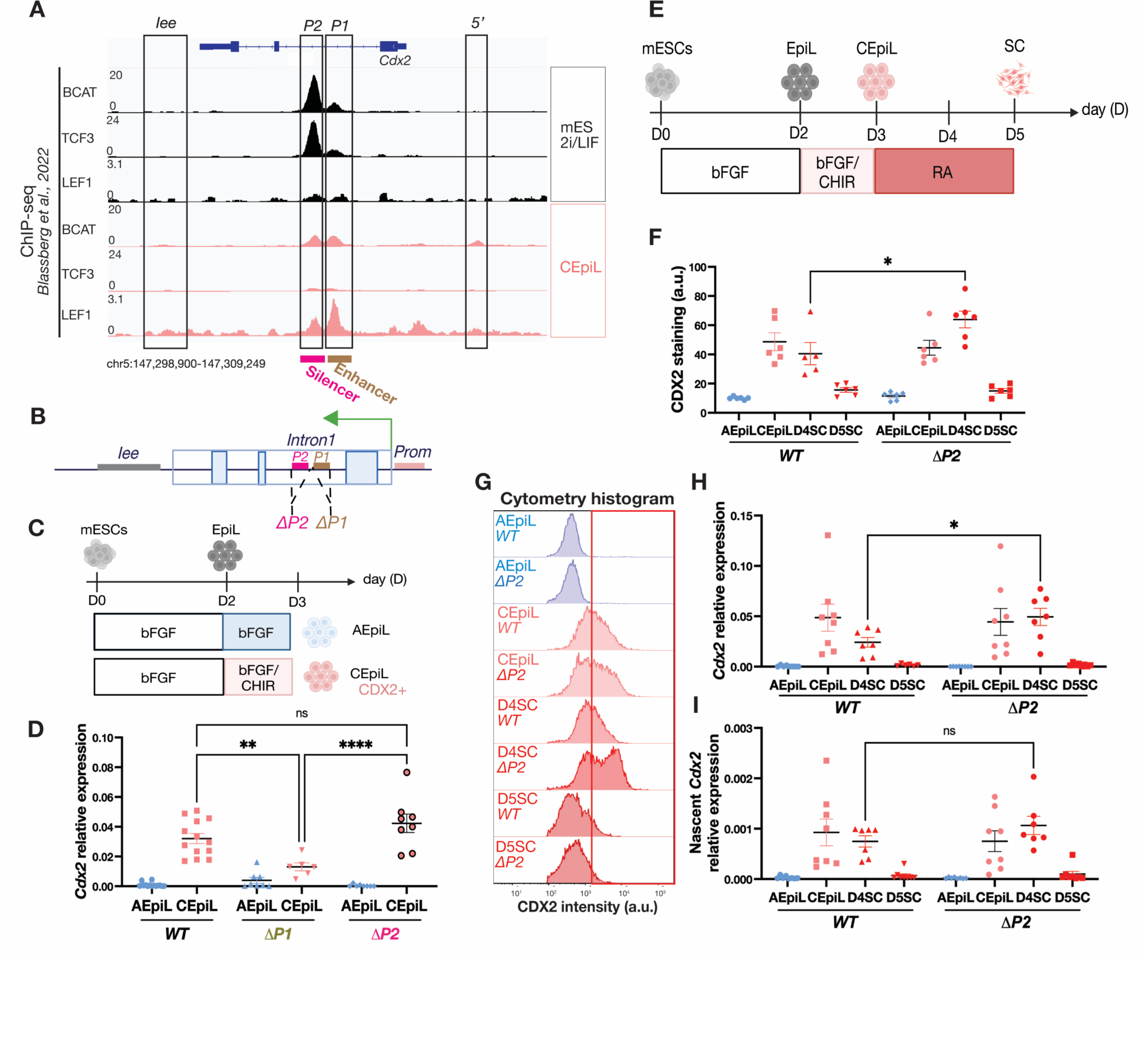
Removal of an intronic silencer extends the duration of *Cdx2* expression in spinal cord progenitors. **(A)** ChIP-seq signal for CTNNB1 (βCAT), TCF3 and LEF1 reveals occupancy at two positions, *P1* and *P2*, within the *Intron1* element of *Cdx2* in naïve ES (black) and CEpiL cells (pink) from 54. (**B**) Schematic of the elements targeted for removal in mouse ESCs corresponding to either *P1* (*ΔP1*) or *P2* (*ΔP2*). (**C**) Conditions used to generate caudal epiblast-like (CEpiL) cells *in vitro* that express *Cdx2* (pink) versus anterior epiblast-like (AEpiL) cells used as a control. (**D**) Relative expression (RT-qPCR) for *Cdx2* in AEpiL (blue) versus CEpiL (pink) conditions from collected from *ΔP1* (n=6), *ΔP2* (n=8) and WT (n=13) cells demonstrates that *Cdx2* is induced in CEpiL cells that lack *P2* (p=0.1592, Tukey’s test) but not in cells lacking *P1* (p=0.0017; Tukey’s test). (**E**) Schematic of the conditions used to differentiate spinal cord (SC) progenitors over a 5-day period (see methods). (**F**) CDX2 levels assessed by flow cytometry in AEpiL (blue), CEpiL (pink) and D4-D5 SC (red) conditions collected from WT (n=5) and *Δ*P2 (n=6) cell lines show increased expression of CDX2 in *ΔP2* D4SC cells compared to WT D4SC cells (p=0.0333, two-tailed t-test). (**G**) Representative cytometry histogram for CDX2 in D3 AEpiL (blue), D3 CEpiL (pink) and D4-D5 spinal cord (red) conditions collected from WT and *ΔP2* cells demonstrates a population of CDX2 positive cells in D4 SC *ΔP2* cells that are not detected in WT D4 SC. (**H-I**) Relative expression (RT-qPCR) for *Cdx2 (H, n=7)* and nascent *Cdx2* (I, n=6) in AEpiL (blue), CEpiL (pink) and D4/D5SC (red) conditions collected from *ΔP2* and WT cells shows significant increased expression of *Cdx2* in *ΔP2* D4SC cells (p=0.0251 for *Cdx2* and p=0.1015 for nascent *Cdx2*; two-tailed t-test). Panels B, C, and E created with BioRender.com.

To test this hypothesis, we generated ESCs lacking either *P1* or *P2* (Fig3B) and directed their differentiation into CEpiL cells as performed above (Fig3C). ESCs lacking *P1*, a region encompassing ∼225 bp, recapitulated the effect of removing the entire intronic CRE (990 bp): *Cdx2* induction was impaired despite exposure to active WNT signalling conditions (Fig3D). By contrast, in the absence of *P2*, cells maintain the ability to induce *Cdx2* (Fig3D). Strikingly, in SC conditions (Fig3E), *P2*-lacking ESCs prolong CDX2 expression, with higher levels of protein (Fig3F-G), spliced transcript (Fig3H) but not nascent transcript (Fig3I) at day 4. In summary, the data indicate that defined subregions within a single intron mediate opposing regulatory outcomes on *Cdx2*, and identify *P2* as a silencer controlling *Cdx2* during the CEpiL to D4SC transition.

### Retinoic acid nuclear receptor composition dictates silencer activity

Having identified two adjacent regions harbouring similar TCF/LEF motifs (Table S1 with functionally opposite regulatory outcomes on *Cdx2*, we next asked what factors recruited to *P1* and *P2* could explain their functional differences. We performed motif analysis to predict transcription factors (TFs) occupying *P1* and *P2* (Table S1 and see Methods). From this analysis, we recovered SOX and TCF/LEF sites at both *P1* and *P2*, consistent with their known occupancy at these sites (Fig3A; FigS3C)^54^. In addition, we detected a striking difference in the composition of RAR versus RXR retinoic acid response elements (RAREs), recognized by the RA family of nuclear receptor TFs (Fig4A and FigS3A-B). Within *P1*, three separate RXRG motifs were detected (Table S1 and Fig4A). By contrast, *P2* lacked any recognizable RXRG motifs, and instead, contained a single RARB motif. Similarly, no occurrences of the RARB site in *P2* were observed in *P1* (Table S1 and Fig4A). In contrast to *Rarg,* which is detected throughout the differentiation, *Rxrg* levels drop while *Rarb* levels rise in single cells as they progress from a CEpiL to SC identity (Fig4B)^24^. This raises the possibility that the relative abundance of nuclear receptors in different cell types ensures context-specific regulatory control over the duration of *Cdx2* via *P1* and *P2*. Consistent with this view, CEpiL cells treated with increasing amounts of all-*trans*-RA (ATRA) display increased levels of *Rarb* in the resulting SC progenitors (Fig4C), while *Cdx2* is reduced (Fig4D).

**Figure 4:**
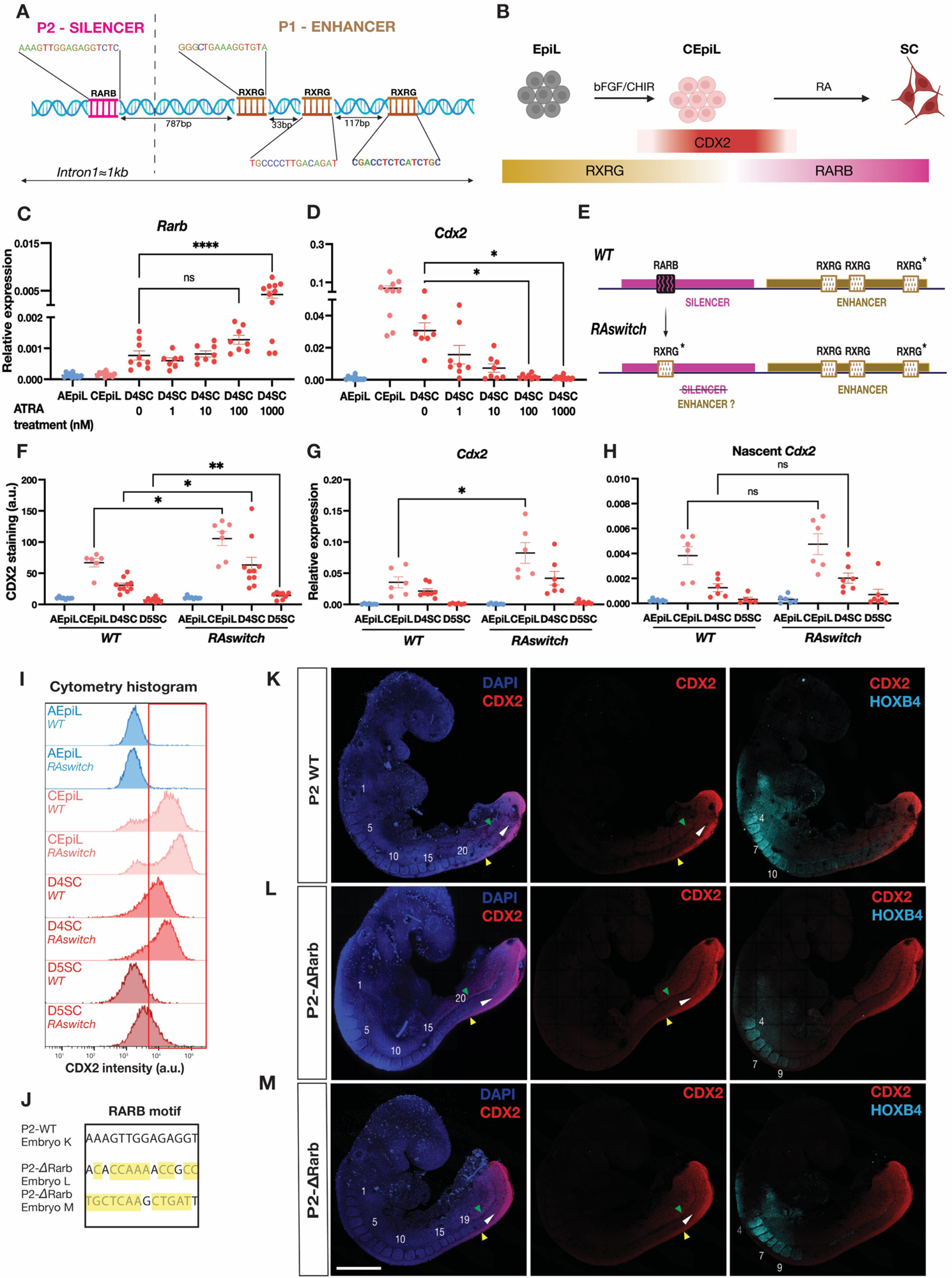
RARE motif switching controls *Cdx2* silencer activity. (**A-B**) Schematic of the RARE binding motifs distributed within the *Intron1* element and the retinoic acid nuclear receptor expression changes observed as mESC differentiate into spinal cord progenitors (24). (**C-D**) Relative expression (RT-qPCR) for *Rarb* (C) and *Cdx2* (D) in anterior epiblast-like (AEpiL, blue), caudal epiblast-like (CEpiL, pink) and day 4 spinal cord (D4SC, red) cells exposed to increasing amounts of all-*trans*-retinoic acid (ATRA) between day 3 and day 4 shows an increase in *Rarb* at 1μM coincides with a significant reduction in *Cdx2* in D4 SC cells (n=11) versus no ATRA treatment control D4SC cells (n=8, p<0.0001 for *Rarb*; p= 0.0160 for *Cdx2*; ANOVA test followed by Tukey’s test). (**E**) *RAswitch* ESCs harbour a single copy of RXRG (indicated with an asterix) instead of an RARB motif at *P2*. (**F**) Quantification of CDX2 levels assessed by flow cytometry in AEpiL (blue), CEpiL (pink), D4 SC and D5 SC (red) conditions collected from *WT* and *RAswitch* cells shows a significant increase in *Cdx2* expression levels in *RAswitch* CEpiL (n=7, p= 0.0163, two-tailed t-test), D4 SC (n=10, p=0.0212, two-tailed t-test) and D5 SC (n=8, p=0.0068, two-tailed t-test) cells compared to WT CEpiL (n=6), D4 SC (n=10) and D5 SC (n=8) cells. (**G-H**) Relative expression (RT-qPCR) for *Cdx2* (G) or nascent *Cdx2* (H) in AEpiL (blue), CEpiL (pink), D4 SC and D5 SC (red) conditions collected from *WT* and *RAswitch* cells shows a significant increase in *Cdx2* levels in *RAswitch* CEpiL (n=6) cells compared to WT CEpiL cells (n=6, p=0.0315, two-tailed t-test; n=7 for WT versus *RAswitch* D4 SC cells, p= 0.0823; two-tailed t-test) which is not detected at the nascent transcript level (H). (**I**) Representative histogram for CDX2 assessed by cytometry in AEpiL (blue), CEpiL (pink), D4 SC and D5 SC (red) conditions collected from *WT* and *RAswitch* cells. (**J**) Resulting genotypes in embryos presented in K-M following CRISPR/Cas9 in fertilized zygotes (see methods) highlighting mis-matches to the WT RARB motif. (**K-M**) Wholemount immunofluorescence for CDX2 (red) and HOXB4 (cyan) together with DAPI (blue) across stage-matched embryos harvested at E9 that contain an intact RARB motif (K, WT *Rarb*) versus mutant RARB motifs (L-M, *P2-ΔRarb*). Somites are numbered in white; green, white and yellow arrows indicate the approximate anterior limit of CDX2 expression in the lateral plate mesoderm, presomitic mesoderm and neural tube, respectively. WT *Rarb* embryos display a caudal limit of HOXB4 that extends to somite 10 (K), versus a caudal limit at somite 9 in *P2-ΔRarb* mutants (L-M). Abnormal tailbud morphology is most apparent in the *P2-ΔRarb* mutant presented in M. Scale bar represents 500μm. Panels A, B, and E created with BioRender.com.

Having established that increased levels of *Rarb* correspond to lower levels of *Cdx2* in SC progenitors, we sought to verify whether the silencing activity of *P2* relies on the presence of the RARB motif in the region. We generated ESCs in which the single RARB motif present in *P2* was replaced with a single copy of the RXRG motif, identical in sequence to the highest-affinity RXRG site detected in *P1* (Fig4E; RXRG site labelled with an asterix), to test the hypothesis that motif substitution underpins the functional differences between *P1* (enhancer) and *P2* (silencer). By switching, rather than deleting, a single motif, this allowed us to test the requirement for distinct RARE motifs within *P2*, while otherwise preserving the DNA sequence length and composition of the *P2* silencer element. The resulting “*RAswitch*” ESCs, which harbour an additional RXRG site compared to control cells, resulted in a small but significant increase in *Cdx2* levels in CEpiL cells, detected at both the transcript and protein level (Fig4F-I, in light pink). In addition, as *RAswitch* cells progressed to a spinal cord identity, CDX2 expression was maintained, in contrast to control cells which instead, begin downregulating *Cdx2*, demonstrating the vital role of the RARE subtype at *P2* to ensure the appropriate silencing of *Cdx2* in SC progenitors (Fig4F-I, in red). By contrast, nascent *Cdx2* transcript was not significantly different in WT versus *RAswitch* cells, in both CEpiL or SC conditions (Fig4H). Taken together, the data demonstrate that the total number and composition of RAREs within P2 determine *Cdx2* regulatory element function.

As the onset and termination of *Cdx2* is mediated through the intronic CRE, also bound by several WNT effectors, how do alternative RA nuclear receptors facilitate *Cdx2* transcriptional control (Fig3A)? Tight control over the level of SOX2 in CEpiL cells is required for *Cdx2* expression, and correlates with genome-wide redistribution of several TCF/LEF effectors, both at the *Cdx2* locus and at a genome-wide scale^54^ (FigS3C). To probe potential interactions between RARB or RXRG and LEF1 or SOX2, we used AlphaFold-multimer^58^ (see methods) to perform *in silico* predictions of possible protein-protein complexes (FigS3D). These simulations provide evidence that both RARB and RXRG can interact with LEF1 and SOX2. Taken together, these findings support the hypothesis that a LEF1/SOX2 complex is redistributed by RA nuclear receptors as cells adopt a spinal cord identity (FigS3E).

### P2 operates as a silencer *in vivo*

Having established a model of *Cdx2* regulatory control during posterior body formation *in vitro*, including the position of a silencer, we sought to validate these findings *in vivo*. To this end, we attempted to perturb silencer activity by disrupting the RARB site within *P2* in mouse embryos. We reasoned that if *P2* functions as a silencer for *Cdx2* during posterior body formation, disruptions to the RARB site alone would be expected to transiently prolong *Cdx2* expression, an effect known to perturb tailbud morphology and *Hox* gene expression boundaries in mouse embryos^2,11^. We used CRISPR-Cas9, together with the same gRNA pairs used *in vitro* to remove *P2* (**△***P2*) or to disrupt directly the RARB site in fertilized zygotes, and collected resulting somite-matched transient transgenic embryos at mid-gestation (Table S2). *P2-ΔRarb* CRISPR mutants recovered at ∼E9 harboured mutations and deletions within the *P2* region (Fig4J). The RARB site was disrupted in mutants (Fig4L-M,J), relative to control littermates that harboured an entirely intact *P2* CRE (*WT P2*; Fig4K,J).

Whole embryo immunofluorescence and imaging confirmed that P2-*WT* embryos recapitulated the known endogenous expression pattern of CDX2 (3,9), which is restricted to the tailbud and the caudal most aspect of the neural tube at this stage (Fig4K). Strikingly, the *P2-ΔRarb* CRISPR mutant embryos displayed a rostral expansion in CDX2, most notably in the mesoderm (Fig4L-M; green and white arrows). In addition, regional identity was disrupted, most notably in the somites. Mutant embryos displayed a caudal limit of HOXB4 that extended to somite 9, in contrast to the control which reached a more caudal position, up to somite 10 (Fig4 compare K to L-M). Mutant embryos also displayed abnormal tailbud morphology, reminiscent of the phenotype resulting from prolonged *Cdx2* expression in mouse embryos^2^. These data provide evidence that *P2* acts as a silencer for *Cdx2*, and pinpoints the RARB motif as a critical site that operates during posterior body formation *in vivo*.

## DISCUSSION

### Multiple, nonredundant CREs coordinate *Cdx2* expression during development

Using an *in vitro* model of posterior body development, we demonstrate that multiple, functionally discrete CREs convert extrinsic cues into a finite window of expression. These data support the idea proposed by previous enhancer reporter experiments that multiple regulatory elements located proximally to the promoter, regulate *Cdx2*^50^. Furthermore, our data extend these findings by demonstrating the functional specificity of CREs during development: *Intron1* is required for induction in CEpiL conditions (Fig1F,I,L), in contrast to the *5*’ CRE which is not required in this context (Fig1G,J,M), but is indispensable at later stages to maintain *Cdx2* transiently in spinal cord or paraxial mesoderm progenitors (Fig2F-I). Previous studies also proposed that a silencer may regulate the caudal expression pattern of *Cdx2*, as individual enhancers demonstrate ectopic reporter activity^50^. Here, we resolve the location of a silencer for *Cdx2*, residing within *Intron1* that we term *P2* and validate its requirements for appropriate CDX2 activity *in vivo*. By contrast, previous studies demonstrate that a fragment containing *P2* displays enhancer activity in transgenic mouse reporter assays^50,56^. Although we cannot exclude the possibility that *P2* displays enhancer activity in an alternative context, and thus may act as a bifunctional element^59–60^, the reported expression pattern of this element displays ectopic activity relative to endogenous *Cdx2*^50^. Our findings highlight that targeted, base-pair substitutions at CREs in their native context can aid in the identification of regulatory regions that include silencer elements. The data support the emerging view that dual enhancer-silencer elements play a fundamental role in gene expression^59,61^.

Our results confirm that individual CREs perform indispensable roles since single CRE deletions are sufficient to perturb the expression window and cannot be compensated for by the presence of alternative, and accessible, CREs (Fig1F,I,L; Fig2F-I; Fig3D-I). The requirement for several, functionally-distinct CREs may ensure robustness in gene expression^62–64^. Consistent with this view, removal of the *5’* or *P2* region within the intronic CRE has a clear but limited effect on *Cdx2* expression, potentially due to the presence of additional CREs that are yet to be resolved. In addition, *P2* extends the expression window in SC progenitors, yet, *Cdx2* is eventually extinguished in these cells (Fig3F-I). The presence of additional, potentially long-range CREs likely explains this effect, and might also buffer fluctuations of extrinsic signalling as observed in the zebrafish neural plate border for *Zic3* expression^66^ and more recently in mouse embryos^64–65^.

### Motif composition dictates silencer function

CDX2 is expressed in response to active WNT signalling conditions and ChIP-seq against LEF1 demonstrates its preferential accumulation at *P1* (Fig3A). However, TCF/LEF binding motifs are also present in multiple *Cdx2* CREs, yet, despite their accessibility, these sites do not compensate for the intronic CRE upon its removal in *ΔIntron1* cells. These findings suggests that chromatin accessibility is not sufficient to predict enhancer function at CREs^55^, which indicates that an additional mechanism is involved.

Recent findings indicate that the specificity of TCF/LEF binding in the genome is driven by context-specific TFs^67^, and, their level of expression^54^. In CEpiL cells, SOX2 levels dictate the genome-wide occupancy of several WNT effectors, including the occupancy of LEF1 and CTNNB1 at the *Cdx2* intronic CRE^54^. Moreover, the activity of the intronic CRE requires SOX2 binding sites^54^, consistent with the view that the recruitment of WNT effectors is driven by cooperation between cell identity-specific TFs. As *P1* and *P2* both harbour SOX2 binding sites and can be occupied by SOX2^54^, an additional molecular determinant must govern *P1* vs *P2* function.

In this study, we provide evidence that silencer function is driven by motif composition. We demonstrate that silencer *P2* can be converted into an enhancer through a single motif switch from RARB to RXRG (Fig4E-I), designed to reconstitute the silencer with a motif normally restricted to the adjacent *P1* enhancer. In contrast to enhancers, relatively few silencers have been identified and functionally validated during development, especially in mammals^68^. As functional validation of repressive elements remains challenging to perform at scale, the mechanisms that distinguish silencer versus enhancer function remain to be elucidated. Nonetheless, dissection of individual elements in different species demonstrates that motif composition can dictate silencer function in *Drosophila*^69–70^. In mammalian genomes, the same CRE can operate as an enhancer or silencer, a function that changes depending on the cellular context^71–75^. Our results demonstrate that a single nuclear receptor motif switch is sufficient to change the function of a given CRE without altering the cellular conditions. These findings support the view that TF engagement explains the versatility of CREs in different cell types^55,74–75^.

### Silencing mechanisms during posterior body development

Although RA is a known major determinant of posterior body formation^3,24,27,35–36, 76–78^, its mechanism of action is not fully understood. Among the predicted 14 000 potential RAREs in the mouse genome^79^, only a handful have been experimentally validated. These include two RARE-containing enhancers responsible for *Hoxa1* expression in the hindbrain^80^; a single atypical RARE required for expression of *Cdx1* in the spinal cord^81–82^ and an RARE in a silencer for *Fgf8* that functions in the developing trunk^83^. Our *in vitro* study identifies one RA-responsive silencer for *Cdx2*, and we provide evidence that it regulates *Cdx2 in vivo,* in line with previous predictions that *Cdx2* may be silenced via an RARE-dependent mechanism^84^. Whether other silencers might be identified based on their sequence similarity to *P2* remains unclear, however, it is unlikely given the atypical and low-affinity RARE sequence identified at *P2* (FigS3A-B) and operating at other RA-responsive genes such as *Cdx1*^82^. In agreement with this, scanning the whole genome for identical sequence motif matches to *P2* only revealed a single match at the first intron of *Kazn* (unpublished observations). Our findings reiterate the necessity for targeted functional validation of predicted regions to resolve CRE function during development.

Our findings also suggest that nuclear receptor composition at CREs might underpin the pleiotropic role of RA during development^85^. Although enhancer *P1* and silencer *P2* appear to recruit similar TFs (SOX2/LEF1; FigS3C^54^), our *in silico* data suggest that RARB and RXRG can both interact with SOX2 (FigS3D), a known determinant of TCF/LEF occupancy at these sites (FigS3C^54^). The RAR nuclear receptor expression difference between caudal epiblast and spinal cord progenitors^24,35,86–88^, and the presence of distinct RARE motifs in opposing functional elements suggest that RARB and RXRG exert distinct roles via these discrete elements. At the *3’* RARE site within the *Hoxa1* locus, RARG recruits the transcriptional co-activators pCIP/p300; however, RA signalling abolishes p300 and instead promotes RARB occupancy and the recruitment of SUZ12^89^. In addition to polycomb-mediated gene silencing, and chromatin compaction^90–91^, HDAC1 recruitment may promote transcriptional silencing, as demonstrated for *Fgf8* in response to RA signalling during posterior body formation^83^. Thus, multiple modes of repression could explain silencer function, and will be addressed by future studies.

Since previously predicted^68,93^ and experimentally validated^92,94–95^ silencers are located relatively close to the TSS of genes and can be found adjacent to enhancers^92–93^, short-range gene silencing mechanisms may represent a more general principle of gene regulation during development. Consistent with this view, *Cdx1* is regulated by a silencer located ∼400bp upstream of the promoter and ∼500bp away from an enhancer^92^. Physical obstruction of individual enhancers or their interaction with the promoter could silence gene expression^39^. CDX factors display a graded expression profile along the rostrocaudal axis^19^, which in turn, plays a central role in constraining regional identity through the regulation of *Hox* genes^2^. That CDX factors contain conserved regulatory elements and play a caudalising role in several species^8,96–100^ suggests that the regulatory principles governing their transient expression may underpin body plan organisation across multiple bilaterian animals.

## Acknowledgements

We thank James Briscoe, Joaquina Delas, Matthias Merkenschlager, Teresa Rayon, Kate Storey, and all lab members for comments on the manuscript. We are grateful to Tristan Rodriguez for advice on the generation of transgenic embryos. For support, training and access to equipment we thank James Elliot from the LMS/NIHR Imperial Biomedical Research Centre Flow Cytometry Facility, the staff at the Central Biomedical Services unit at Imperial College London, Zoe Webster at the LMS transgenic facility, and Dirk Dormann at the LMS microscopy facility. This work was supported by a Sir Henry Dale Fellowship awarded to VM, jointly funded by the Wellcome Trust and the Royal Society (Grant Number 218536/Z/19/Z).

## Author contributions

IA and VM conceived the project, designed the experiments, interpreted the data, and wrote the manuscript. IA performed the experiments and data analysis. DB performed data analysis and together with BL interpreted the data. BM performed microinjections and embryo transfers. All authors revised the manuscript.

## METHODS

### Cells lines

All mouse ESC lines were cultured at 37 °C with 5% CO_2_. All ESC lines used were derived from the XY HM1 line^101^, which was used as the WT control. *ΔIntron1*, *Δ5’*, *ΔIee*, *ΔProm*, *ΔTee*, *ΔP1* and *ΔP2* lines were generated by electroporating pairs of CRISPR targeted to both extremities of the regions of interest (Table S3). After puromycin selection (at a concentration of 1.5 ug/mL), 10 clones were picked and expanded. gDNA was extracted using the PureLink kit (PureLink™ Genomic DNA) according to the manufacturer’s instructions. Clones were then genotyped (Table S3), validated by DNA sequencing, and routinely tested for mycoplasma. For most of the generated KO cell lines, we obtained similar results with a second clone (*ΔIntron1*, *ΔTee*, *ΔProm*, *ΔP2*, *ΔIee*).

The *RAswitch* line was created using HDR recombinant oligos electroporated with the sgRNA guide (Table S3) and the Cas9 protein (#1081058) supplemented with the Alt-R Cas9 electroporation enhancer (#1075915) into HM1 cells. After recovery using the Alt-R HDR EnhancerV2 (#10007910), 10 clones were picked, expanded, and, genotyped (Table S3) and validated by DNA sequencing in a similar method as described above.

### ESC culture and differentiation

All mouse ESCs were expanded on mitotically inactivated mouse embryonic fibroblasts (feeders) in ESC medium (DMEM knockout medium supplemented with 1.000U/ml LIF, 10% cell-culture-validated foetal bovine serum, and 2mM L-Glutamine).

To differentiate mESCs into neural or paraxial mesoderm progenitors, ESCs were differentiated as previously described^52,24^. Briefly, ESCs were dissociated with 0.05% trypsin, and plated on tissue-culture-treated plates for two sequential 20 minutes (mins) periods in ESC medium to separate them from their feeder layer cells, which adhere to the plastic. To start the differentiation, cells remaining in the supernatant were pelleted by centrifugation, counted, and resuspended in N2B27 medium containing 10 ng/ml bFGF, and 40 000 cells per 35 mm gelatin-coated CellBIND dish or 6-well plate (Corning) were plated. N2B27 medium contained a 1:1 ratio of DMEM/F12:Neurobasal medium (Gibco) supplemented with 0.5% N2 (Gibco), 1% B27 (Gibco), 2mM L-glutamine (Gibco), 40mg/ml BSA (Sigma), and 0.1mM 2-mercaptoethanol.

To generate Epiblast-like (EpiL) or Anterior Epiblast-like (AEpiL) cells, the cells were grown respectively for 2 and 3 days in N2B27 +10 ng/ml bFGF. To generate Caudal Epiblast-like (CEpiL) cells, cells were cultured with N2B27 + 10 ng/ml bFGF for 2 days, then N2B27 + 10 ng/ml bFGF +5 μM CHIR99021 for a further day. CEpiL cells were differentiated to spinal cord neural progenitors by continuing the differentiation up to day 5 in N2B27 media containing 10nM all-*trans*-retinoic acid (ATRA, 10nM). In the experiments presented in Fig4C-D, CEpiL were either exposed to N2B27 alone for one day or exposed to concentrations of ATRA ranging from 1nM to 1uM.

To generate paraxial mesoderm progenitors, CEpiL cells (generated as described above) were exposed to 5μM CHIR for a further 2 days. For the paraxial mesoderm differentiation, a different N2B27 media was applied to CEpiL by using B27 devoid of Vitamin A as previously described^24^. Media was refreshed every day from day 2-5 for all experiments.

Details of key compounds are provided in Table S4.

### Immunofluorescence on cells

Cells were washed in PBS and fixed in 4% paraformaldehyde in PBS for 30 mins at 4 °C, followed by three washes in PBS. Primary antibodies (Table S5) were applied overnight at 4°C diluted in filtered blocking solution (2% BSA diluted in PBST – 0.1% Triton X-100 diluted in PBS). Cells were washed for 5 mins three times in PBST and incubated with secondary antibodies (Table S5) at room temperature, for 90 mins. Cells were washed for 5 mins three times in PBST, incubated with DAPI for 15 mins in PBS and washed twice before mounting with a glass coverslip using Prolong Gold (Invitrogen) or kept in PBS for further imaging. Cells were imaged on an inverted SP5 or upright SP5 II confocal microscope (Leica). Z stacks were acquired using the Leica LAS AF software and represented as maximum intensity projections using ImageJ software. The same settings were applied to all images. Images presented in Fig1L-N; FigS1E-F; are representative images of a minimum of three biological replicates.

### Flow cytometry

Cells were washed in PBS and dissociated with Accutase (Gibco). Once detached, cells were collected, washed with PBS, and pelleted. Cells were resuspended in PBS supplemented with live dye (1/1000, Thermo Fisher) and kept in dark at 4° for 30 mins. Cells were pelleted, washed in PBS, pelleted, and resuspended in 4% paraformaldehyde in PBS. Following 15 mins incubation at 4°C, cells were centrifuged, resuspended in PBS, and stored at 4°C for future analysis.

On the day of flow cytometry, cells were transferred for staining in U-bottom 96-well plates. Samples were pelleted and resuspended in 50μl block media (2% BSA diluted in PBST). After 30 mins incubation at room temperature in the platform rocker, antibodies were added to the sample and incubated overnight at 4°C on a platform rocker. Details of primary and secondary antibodies are described in Table S5. Cells were pelleted for 4 mins, washed in PBST, pelleted, and incubated in 50μl PBST supplemented with secondary antibodies (concentration: 1/500) in the dark for 2h at room temperature in the platform rocker. One additional wash was performed before acquisition on a SymphonyA3 (BD Biosciences) using FACSDiva. Analysis was performed using FlowJo.

### RNA extraction, cDNA synthesis and RT-qPCR analysis

RNA used for real time quantitative PCR (RT-qPCR) was extracted from cells using a QIAGEN RNeasy kit in RLT buffer, following the manufacturer’s instructions. Extracts were digested with DNase I to eliminate genomic DNA.

First-strand cDNA synthesis was performed using Superscript III (Invitrogen) using random hexamers and was amplified using PowerUp SYBR-Green Mastermix (Applied Biosystems). RT-qPCR was performed using the Applied Biosystems QuantStudio Real Time PCR system and analysed with Applied Biosystems QuantStudio 12K Flex software. PCR primers were designed using the online PrimerBLAST design tool and validated (standard curve and melting curve) or taken from previously published papers. Primer sequences are detailed in Table S6. Two technical replicates were obtained for each sample and averaged before normalization and statistical analysis. Relative expression values for each gene were calculated by normalization against β-actin, using the delta–delta CT method. RT-qPCR analysis was performed on samples obtained from a minimum of three independent experiments for every primer pair analysed.

### Generation of CRISPR mutant embryos

For generation of the *P2-DRarb* CRISPR mutants, gRNA sequences (Table S2) were ordered as oligonucleotides (Integrated DNA Technologies) together with recombinant Cas9 protein (Alt-R™ S.p. Cas9 Nuclease V3; 1081058). The sgRNA (at 25 ng/µl) and the Cas9 (at 75 ng/µl) were combined and microinjected into the pronuclei of one-cell embryos in two separate rounds of injection (Fig4).

One-cell embryos obtained by super-ovulating 10 C57Bl/6 females with 50 IU of PMSg 48h hours before mating and with 50 IU of HCG on the day of mating were mated with C57Bl/6 stud males. 24h after mating embryos were harvested, cleaned and placed in culture media (KSOM) at 37°C. Each zygote was then microinjected into the pronuclei with the CRISPR/Cas9 complex. Microinjected zygotes were transferred back into recipient females (B6CBAF1; previously mated and plugged by vasectomised CD1 males) by embryo transfer procedure at 0.5dpc. All females were monitored daily in a Biological Support Unit. The recipient females were humanely killed and embryos were harvested at 9.5dpc.

After dissection and collection of amniotic tissue for genotyping, embryos were fixed in 4% paraformaldehyde in PBS for 90 mins at 4°C under gentle agitation, followed by two washes in PBS.

Embryos were genotyped using the HotSHOT DNA (102) extraction protocol. Briefly, amnions were incubated at 95°C for 30 mins in 25uL alkaline lysis buffer before addition of 25uL neutralizing buffer and storage at 4°C. The *Intron1* fragment was amplified by PCR using the primers used for the genotyping of the *ΔIntron1* cell line (provided Table S3).

After purification using the Qiagen PCR purification kit, the fragment was sequenced using the primer outlined in Table S3.

All animal procedures were performed by certified staff in the Imperial College London Central Biomedical Services Facility in accordance with the Animal (Scientific Procedures) Act 1986 under the UK Home Office project license PP2904879.

### Embryo wholemount immunofluorescence

Embryos were permeabilized in 0.5% Triton X-100 diluted in PBS for 30 mins at room temperature under gentle agitation. After permeabilization, embryos were incubated in filtered block media (2% BSA and 4% donkey serum diluted in PBST) at room temperature for 2 hrs under gentle agitation. Primary antibodies (Table S5) were applied overnight at 4 °C diluted in filtered block media under gentle agitation. The following morning, embryos were washed for 2 hrs in PBST, 4-5 times, at room temperature under gentle agitation and incubated in filtered block media overnight at 4°C under gentle agitation. Secondary antibodies (Table S5) were applied diluted in PBST (1/500) at room temperature for 90 mins in the dark under gentle agitation. After 10 mins PBST washes, embryos were incubated with DAPI (1/1000) in PBST in the dark under gentle agitation at room temperature for 30 mins.

After PBST washes, embryos were mounted in 1.5% low-melt agarose in p35 Ibidi plates before imaging using an inverted Leica DLS. Z stacks were acquired using the Leica LAS AF software and represented as maximum intensity projections using ImageJ software.

### ChIP-seq, ATAC-seq and mRNA-seq data and processing

ATAC-seq data from day 2 epiblast-like (D2-EpiL), day 3 caudal epiblast-like (D3-CEpiL), day 4 (D4-) and day 5 (D5-) spinal cord (SC) were obtained from Metzis et al., 2018^52^ (accession number E-MTAB-6337, Table S7). Pseudo-bulk ATAC-seq generated from mouse embryo 10x multiome experiments were obtained from Argelaguet et al., 2022^53^ (accession number GSE205117, Table S7). ChIP-seq data from naïve mouse ESCs, caudal epiblast-like cells and *Sox2* over-expressing caudal-epiblast-like cells (Fig3A and FigS3C) were obtained from Blassberg et al., 2022^54^ (accession number GSE162774, Table S7). Big Wig tracks were visualised using IGV^103^.

For mRNA-seq data (Table S7), the nf-core/rnaseq pipeline (version 2.0)^104^ was used with default parameters. Briefly, the pipeline performs quality control, trimming (using TrimGalore!), (pseudo-)alignment (using Salmon), and produces a gene expression matrix. All data were processed relative to the mouse UCSC mm10 genome (UCSC) downloaded from AWS iGenomes (https://github.com/ewels/AWS-iGenomes).

### Identification of Transcription Factor Binding Sites (TFBS)

To identify transcription factor binding sites (TFBS) in genomic sequences, we used a comprehensive bioinformatics approach with publicly available databases and specialised software tools. The matrices representing the binding preferences of transcription factors were obtained from the JASPAR2022 database^105^.

To detect TFBS instances within our specific sequences, we used TFBStools version 1.40.0, as described in ^106^. We set TFBStools to search for matches to the JASPAR matrices within our sequences, specifying an identity match threshold of 80%. The threshold was selected to identify low-affinity sites for TFs of interest in our specified regions. Then, we filtered our list to remove any non-expressed transcription factors based on mRNA-seq data generated *in vitro* in EpiL, CEpiL, D4 SC and D5 SC^31^.

The GERP score from Ensembl, presented FigS3A-B, and calculated from 91 mammalian genomes, was used to estimate the level of conservation of the TFBS (http://ftp.ensembl.org/pub/release-111/compara/conservation_scores/91_mammals.gerp_conservation_score/gerp_conservation_scores.mus_musculus.GRCm39.bw).

### Prediction of Protein-Protein interaction Complexes

To further investigate the functional implications of identified TFBS instances, we aimed to predict the structures of protein-protein complexes involving our TFs of interest. To do this, we retrieved the complete TF protein sequences from InterPro, a comprehensive database of protein families, domains, and functional sites^108^. We then used ColabFold version 1.3.0, which is an interface for AlphaFold-multimer program^107^. AlphaFold-multimer is a state-of-the-art method for predicting protein complex structures, leveraging deep learning to estimate the three-dimensional arrangements of protein subunits^58,109^. By inputting the TF protein sequences into ColabFold, we were able to obtain high-confidence predictions of their potential interactions and complex formations.

### Statistics and reproducibility

No statistical method was used to pre-determine sample size. No data were excluded from the analyses. The experiments were not randomized. The investigators were not blinded to allocation during experiments and outcome assessment. For all statistical analyses, data were obtained from a minimum of three independent experiments. Bars denote mean +/- s.e.m and statistical significance was calculated using GraphPad Prism (GraphPad Software).

Details of replicate numbers, and statistics for each experiment are specified in the figure legends.

---

**Figure S1.**
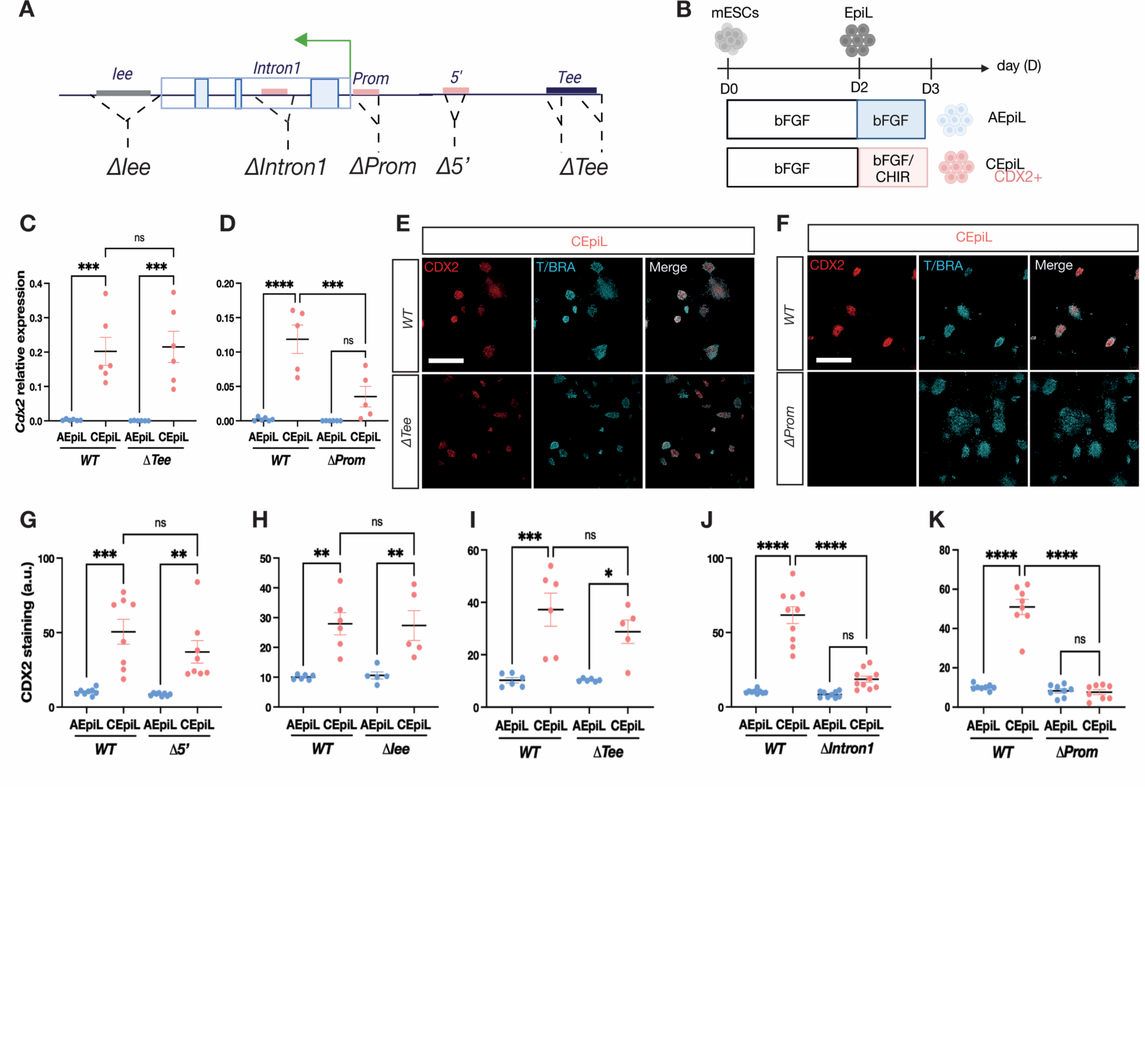
*Cdx2* expression in caudal epiblast progenitors is not affected by the removal of the *Tee* or *Iee* element. Related to Figure 1. **(A)** Schematic of the *Cdx2* locus illustrating the position of elements targeted for removal in mouse ESCs (see methods). (**B**) Conditions used to generate caudal epiblast-like (CEpiL) cells *in vitro* that express *Cdx2* (pink), versus anterior epiblast-like (AEpiL) cells used as a control (blue). (**C-D**) Relative expression (RT-qPCR) for *Cdx2* in AEpiL (blue) versus CEpiL (pink) conditions from collected from cells lacking the TEE element (*ΔTee*, C, n=6), or the promoter (*ΔProm,* D, n=6) demonstrates that *Cdx2* is lost in CEpiL cells lacking the promoter (D, p=0.0006, Tukey’s test) but not *Tee*-lacking cells (p=0.9898; Tukey’s test). (**E-F**) CDX2 (red) and T/BRA (cyan) immunofluorescence for WT and *ΔProm* (F) CEpiL cells confirms a loss of CDX2 in promoter-lacking but not *ΔTee* cells (E); scale bar represents 500μm. n=3. (**G-K**) CDX2 levels assessed by cytometry in AEpiL (blue) versus CEpiL (pink) conditions collected from *Δ5’* (G, n=8), *ΔIee* (H, n=5), *ΔTee* (I, n=6), *ΔIntron1* (J, n=10) and *ΔProm* (K, n=8) cells shows CDX2 is not induced in CEpiL cells that lack the *Intron1* element (J, p<0.0001, Tukey’s test) or the promoter element (K, p <0.0001, Tukey’s test) while the removal of the *5’* (G, p= 0.3495), *Tee* (I, p= 0.4537) and *Iee* (H, p= 0.9990) has no effect on *Cdx2*. Panels A-B created with BioRender.com.

**Figure S2:**
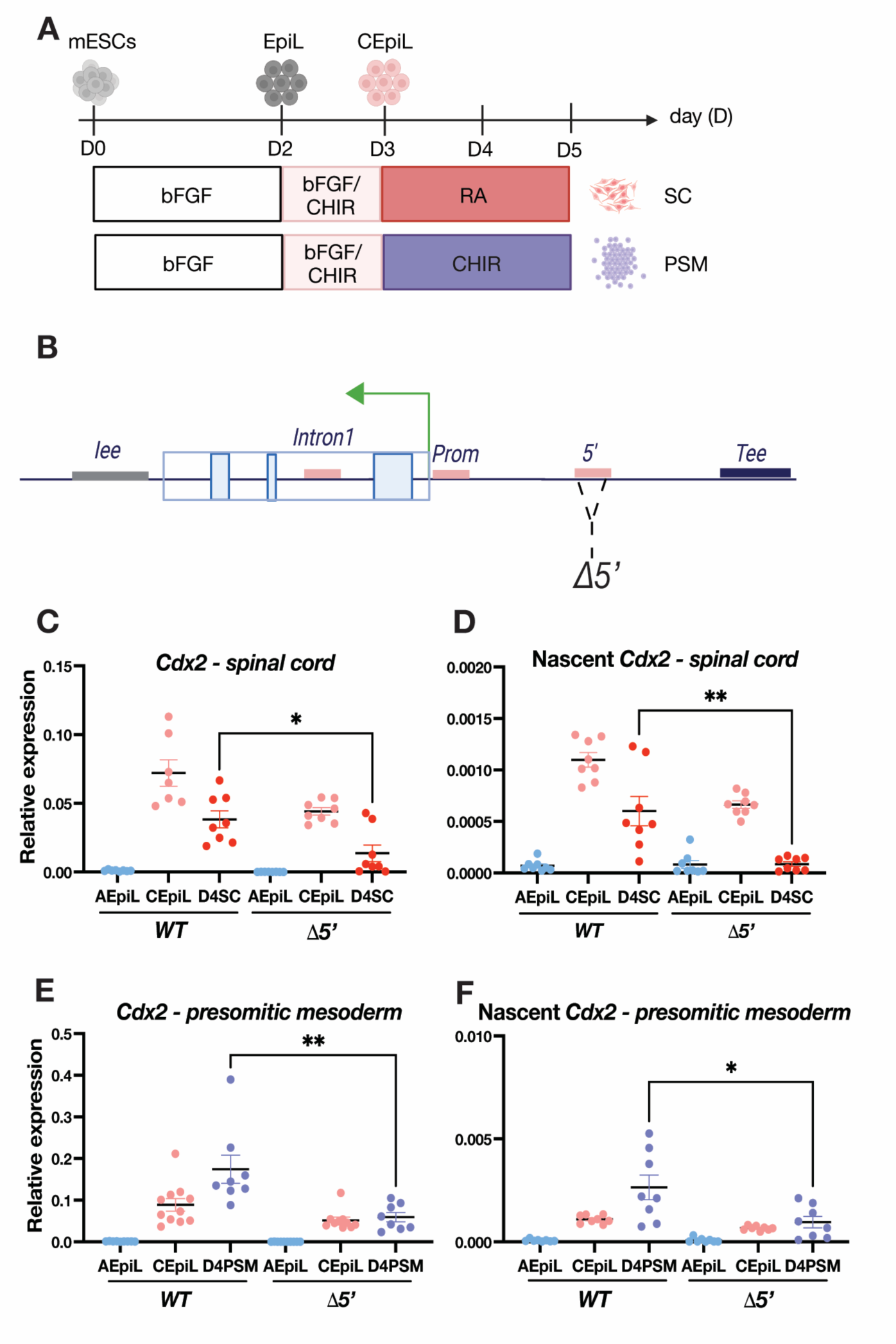
Removal of the 5’ element disrupts *Cdx2* nascent transcription. Related to Figure 2. **(A)** Schematic of the conditions used to generate caudal epiblast like (CEpiL) cells from mouse ESCs for subsequent differentiation into spinal cord (SC) (top row) or presomitic mesoderm (PSM; bottom row) progenitors over a 5-day (D) period (see methods). (**B**) Schematic illustrating the position of the 5’ element targeted for removal in mouse ESCs using pairs of CRISPR (*Δ5’*) (see methods). (**C-F**) Relative expression (RT-qPCR) detecting *Cdx2* spliced (C, E) versus nascent *Cdx2* transcript (D, F) in AEpiL (blue), CEpiL (pink), D4SC (red) and D4 PSM conditions collected from WT and *Δ5*’cells show a statistically significant reduction in *Cdx2* in *Δ5*’ D4 SC cells (C, n=8, p= 0.0129; D, n=8, p=0.0031; two-tailed t-test) versus *Δ5*’ D4PSM cells (E, n=8; p= 0.006; F, n=8; p=0.0233; two-tailed t-test). Panels A-B created with BioRender.com.

**Figure S3:**
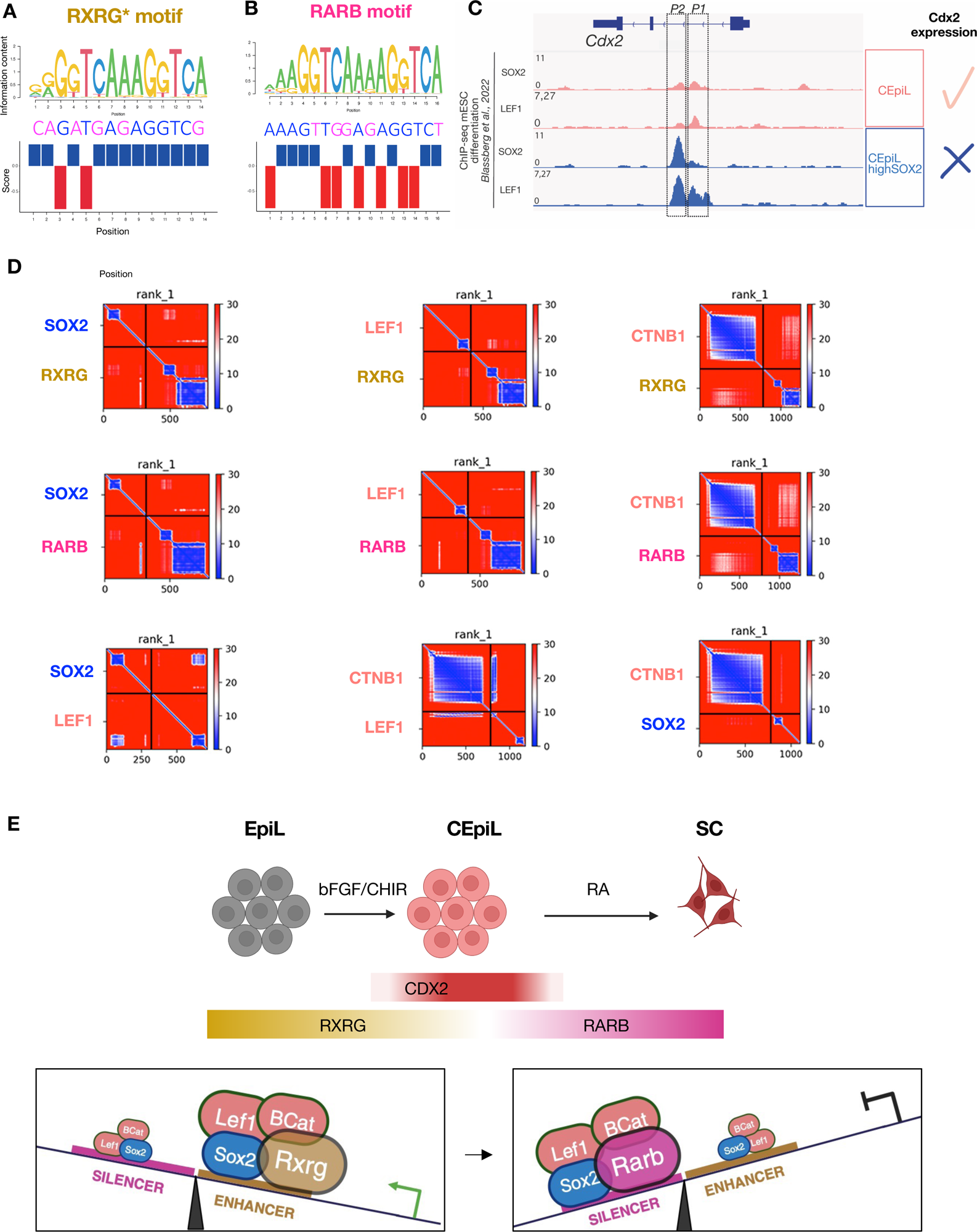
RA and WNT signalling effectors interact in silico. Relates to Figure 3. (**A-B**) Alignment between RARB (A) and RXRG (B) transcription factor binding sites (TFBS) extracted from JASPAR with their respective TFBS matches in *P2* and *P1*. The motif is represented as the sequence logo, showing the nucleotide preference of the motif at each position. Blue and magenta letters in the genomic TFBS sequence represent matches and mismatches with the motif consensus sequence, respectively. The bottom panel shows the TFBS conservation. The positive values mean evolutionary constraints, while positions with negative values are under positive selection. (**C**) ChIP-seq signals at the *Cdx2* locus showing occupancy of SOX2 and LEF1 in CEpiL cells generated *in vitro* from WT caudal epiblast-like cells (CEpiL, in pink) versus CEpiL cells expressing high levels of SOX2 (CEpiL highSOX2, in blue) that repress *Cdx2* (from 54). (**D**) Protein contact probability matrices calculated by AlphaFold-Multimer. Lower values represent higher contact probabilities. Protein sequences are divided by the black line. The upper right and lower left quadrants represent predicted protein-protein interactions. (**E**) Hypothetical model of cell type specific regulation of *Cdx2* driven by alterations in the nuclear receptor expression via enhancer *P1* versus silencer *P2*. In caudal epiblast cells, RXRG is expressed, favouring the recruitment of SOX2/LEF1/CTNB1 containing complexes at *P1* to promote expression of *Cdx2*. By contrast, a rise in retinoic acid signalling results in *Rarb* expression and loss of *Rxrg* in spinal cord progenitors. RARB redistributes SOX2/LEF1/beta-catenin containing complexes to silencer *P2* in SC progenitors. Panel E created with BioRender.com

## REFERENCES

1. Davidson, E.H. (2010). Emerging properties of animal gene regulatory networks. Nature 468, 911–920. 10.1038/nature09645.

2. Gaunt, S.J., Drage, D., and Trubshaw, R.C. (2008). Increased Cdx protein dose effects upon axial patterning in transgenic lines of mice. Development 135, 2511– 2520. 10.1242/dev.015909.

3. Young, T., Rowland, J.E., Van De Ven, C., Bialecka, M., Novoa, A., Carapuco, M., Van Nes, J., De Graaff, W., Duluc, I., Freund, J.-N., et al. (2009). Cdx and Hox Genes Differentially Regulate Posterior Axial Growth in Mammalian Embryos. Developmental Cell 17, 516–526. 10.1016/j.devcel.2009.08.010.

4. Van Rooijen, C., Simmini, S., Bialecka, M., Neijts, R., Van De Ven, C., Beck, F., and Deschamps, J. (2012). Evolutionarily conserved requirement of Cdx for post-occipital tissue emergence. Development 139, 2576–2583. 10.1242/dev.079848.

5. Neijts, R., Amin, S., Van Rooijen, C., and Deschamps, J. (2017). Cdx is crucial for the timing mechanism driving colinear Hox activation and defines a trunk segment in the Hox cluster topology. Developmental Biology 422, 146–154. 10.1016/j.ydbio.2016.12.024.

6. Van Den Akker, E., Forlani, S., Chawengsaksophak, K., De Graaff, W., Beck, F., Meyer, B.I., and Deschamps, J. (2002). *Cdx1* and *Cdx2* have overlapping functions in anteroposterior patterning and posterior axis elongation. Development 129, 2181– 2193. 10.1242/dev.129.9.2181.

7. Savory, J.G.A., Pilon, N., Grainger, S., Sylvestre, J.-R., Béland, M., Houle, M., Oh, K., and Lohnes, D. (2009). Cdx1 and Cdx2 are functionally equivalent in vertebral patterning. Developmental Biology 330, 114–122. 10.1016/j.ydbio.2009.03.016.

8. Chawengsaksophak, K., De Graaff, W., Rossant, J., Deschamps, J., and Beck, F. (2004). *Cdx2* is essential for axial elongation in mouse development. Proc. Natl. Acad. Sci. U.S.A. 101, 7641–7645. 10.1073/pnas.0401654101.

9. Amin, S., Neijts, R., Simmini, S., Van Rooijen, C., Tan, S.C., Kester, L., Van Oudenaarden, A., Creyghton, M.P., and Deschamps, J. (2016). Cdx and T Brachyury Co-activate Growth Signaling in the Embryonic Axial Progenitor Niche. Cell Reports 17, 3165–3177. 10.1016/j.celrep.2016.11.069.

10. Needham, J., and Metzis, V. (2022). Heads or tails: making the spinal cord. Developmental Biology 485, 80–92. 10.1016/j.ydbio.2022.03.002.

11. Charité, J., Graaff, W.D., Consten, D., Reijnen, M.J., Korving, J., and Deschamps, J. (1998). Transducing positional information to the *Hox* genes: critical interaction of *cdx* gene products with position-sensitive regulatory elements. Development 125, 4349–4358. 10.1242/dev.125.22.4349.

12. Bel-Vialar, S., Itasaki, N., and Krumlauf, R. (2002). Initiating Hox gene expression: in the early chick neural tube differential sensitivity to FGF and RA signaling subdivides the *HoxB* genes in two distinct groups. Development 129, 5103–5115. 10.1242/dev.129.22.5103.

13. Shimizu, T., Bae, Y.-K., and Hibi, M. (2006). Cdx-Hox code controls competence for responding to Fgfs and retinoic acid in zebrafish neural tissue. Development 133, 4709–4719. 10.1242/dev.02660.

14. Skromne, I., Thorsen, D., Hale, M., Prince, V.E., and Ho, R.K. (2007). Repression of the hindbrain developmental program by Cdx factors is required for the specification of the vertebrate spinal cord. Development 134, 2147–2158. 10.1242/dev.002980.

15. Sturgeon, K., Kaneko, T., Biemann, M., Gauthier, A., Chawengsaksophak, K., and Cordes, S.P. (2011). Cdx1 refines positional identity of the vertebrate hindbrain by directly repressing *Mafb* expression. Development 138, 65–74. 10.1242/dev.058727.

16. Van Nes, J., De Graaff, W., Lebrin, F., Gerhard, M., Beck, F., and Deschamps, J. (2006). The *Cdx4* mutation affects axial development and reveals an essential role of Cdx genes in the ontogenesis of the placental labyrinth in mice. Development 133, 419–428. 10.1242/dev.02216.

17. Young, T., and Deschamps, J. (2009). Chapter 8 Hox, Cdx, and Anteroposterior Patterning in the Mouse Embryo. In Current Topics in Developmental Biology (Academic Press), pp. 235–255. 10.1016/S0070-2153(09)88008-3.

18. Beck, F., Erler, T., Russell, A., and James, R. (1995). Expression of Cdx-2 in the mouse embryo and placenta: Possible role in patterning of the extra-embryonic membranes. Dev. Dyn. 204, 219–227. 10.1002/aja.1002040302.

19. Gaunt, S.J., Drage, D., and Trubshaw, R.C. (2005). cdx4/lacZ and cdx2/lacZ protein gradients formed by decay during gastrulation in the mouse. Int. J. Dev. Biol. 49, 901–908. 10.1387/ijdb.052021sg.

20. McDole, K., and Zheng, Y. (2012). Generation and live imaging of an endogenous Cdx2 reporter mouse line. Genesis 50, 775–782. 10.1002/dvg.22049.

21. Tzouanacou, E., Wegener, A., Wymeersch, F.J., Wilson, V., and Nicolas, J.-F. (2009). Redefining the Progression of Lineage Segregations during Mammalian Embryogenesis by Clonal Analysis. Developmental Cell 17, 365–376. 10.1016/j.devcel.2009.08.002.

22. Henrique, D., Abranches, E., Verrier, L., and Storey, K.G. (2015). Neuromesodermal progenitors and the making of the spinal cord. Development 142, 2864–2875. 10.1242/dev.119768.

23. Binagui-Casas, A., Dias, A., Guillot, C., Metzis, V., and Saunders, D. (2021). Building consensus in neuromesodermal research: Current advances and future biomedical perspectives. Current Opinion in Cell Biology 73, 133–140. 10.1016/j.ceb.2021.08.003.

24. Gouti, M., Delile, J., Stamataki, D., Wymeersch, F.J., Huang, Y., Kleinjung, J., Wilson, V., and Briscoe, J. (2017). A Gene Regulatory Network Balances Neural and Mesoderm Specification during Vertebrate Trunk Development. Developmental Cell 41, 243–261.e7. 10.1016/j.devcel.2017.04.002.

25. Guibentif, C., Griffiths, J.A., Imaz-Rosshandler, I., Ghazanfar, S., Nichols, J., Wilson, V., Göttgens, B., and Marioni, J.C. (2021). Diverse Routes toward Early Somites in the Mouse Embryo. Developmental Cell 56, 141–153.e6. 10.1016/j.devcel.2020.11.013.

26. Pijuan-Sala, B., Griffiths, J.A., Guibentif, C., Hiscock, T.W., Jawaid, W., Calero-Nieto, F.J., Mulas, C., Ibarra-Soria, X., Tyser, R.C.V., Ho, D.L.L., et al. (2019). A single-cell molecular map of mouse gastrulation and early organogenesis. Nature 566, 490–495. 10.1038/s41586-019-0933-9.

27. Zhao, X., and Duester, G. (2009). Effect of retinoic acid signaling on Wnt/β-catenin and FGF signaling during body axis extension. Gene Expression Patterns 9, 430–435. 10.1016/j.gep.2009.06.003.

28. Ikeya, M., and Takada, S. (2001). Wnt-3a is required for somite specification along the anteroposterior axis of the mouse embryo and for regulation of cdx-1 expression. Mechanisms of Development 103, 27–33.

29. Keenan, I.D., Sharrard, R.M., and Isaacs, H.V. (2006). FGF signal transduction and the regulation of Cdx gene expression. Developmental Biology 299, 478–488. 10.1016/j.ydbio.2006.08.040.

30. Frith, T.J., Granata, I., Wind, M., Stout, E., Thompson, O., Neumann, K., Stavish, D., Heath, P.R., Ortmann, D., Hackland, J.O., et al. (2018). Human axial progenitors generate trunk neural crest cells in vitro. eLife 7, e35786. 10.7554/eLife.35786.

31. Gouti, M., Tsakiridis, A., Wymeersch, F.J., Huang, Y., Kleinjung, J., Wilson, V., and Briscoe, J. (2014). In Vitro Generation of Neuromesodermal Progenitors Reveals Distinct Roles for Wnt Signalling in the Specification of Spinal Cord and Paraxial Mesoderm Identity. PLoS Biol 12, e1001937. 10.1371/journal.pbio.1001937.

32. Lippmann, E.S., Williams, C.E., Ruhl, D.A., Estevez-Silva, M.C., Chapman, E.R., Coon, J.J., and Ashton, R.S. (2015). Deterministic HOX Patterning in Human Pluripotent Stem Cell-Derived Neuroectoderm. Stem Cell Reports 4, 632–644. 10.1016/j.stemcr.2015.02.018.

33. Mazzoni, E.O., Mahony, S., Peljto, M., Patel, T., Thornton, S.R., McCuine, S., Reeder, C., Boyer, L.A., Young, R.A., Gifford, D.K., et al. (2013). Saltatory remodeling of Hox chromatin in response to rostrocaudal patterning signals. Nat Neurosci 16, 1191–1198. 10.1038/nn.3490.

34. Wind, M., Gogolou, A., Manipur, I., Granata, I., Butler, L., Andrews, P.W., Barbaric, I., Ning, K., Guarracino, M.R., Placzek, M., et al. (2021). Defining the signalling determinants of a posterior ventral spinal cord identity in human neuromesodermal progenitor derivatives. Development 148, dev194415. 10.1242/dev.194415.

35. Del Corral, R.D., Olivera-Martinez, I., Goriely, A., Gale, E., Maden, M., and Storey, K. (2003). Opposing FGF and Retinoid Pathways Control Ventral Neural Pattern, Neuronal Differentiation, and Segmentation during Body Axis Extension. Neuron 40, 65–79. 10.1016/S0896-6273(03)00565-8.

36. Olivera-Martinez, I., and Storey, K.G. (2007). Wnt signals provide a timing mechanism for the FGF-retinoid differentiation switch during vertebrate body axis extension. Development 134, 2125–2135. 10.1242/dev.000216.

37. Cooper, F., Gentsch, G.E., Mitter, R., Bouissou, C., Healy, L.E., Rodriguez, A.H., Smith, J.C., and Bernardo, A.S. (2022). Rostrocaudal patterning and neural crest differentiation of human pre-neural spinal cord progenitors in vitro. Stem Cell Reports 17, 894–910. 10.1016/j.stemcr.2022.02.018.

38. Long, H.K., Prescott, S.L., and Wysocka, J. (2016). Ever-Changing Landscapes: Transcriptional Enhancers in Development and Evolution. Cell 167, 1170–1187. 10.1016/j.cell.2016.09.018.

39. Ogbourne, S., and Antalis, T.M. (1998). Transcriptional control and the role of silencers in transcriptional regulation in eukaryotes. Biochemical Journal 331, 1–14. 10.1042/bj3310001.

40. Herold, M., Bartkuhn, M., and Renkawitz, R. (2012). CTCF: insights into insulator function during development. Development 139, 1045–1057. 10.1242/dev.065268.

41. Batut, P.J., Bing, X.Y., Sisco, Z., Raimundo, J., Levo, M., and Levine, M.S. (2022). Genome organization controls transcriptional dynamics during development. Science 375, 566–570. 10.1126/science.abi7178.

42. Blayney, J.W., Francis, H., Rampasekova, A., Camellato, B., Mitchell, L., Stolper, R., Cornell, L., Babbs, C., Boeke, J.D., Higgs, D.R., et al. (2023). Super-enhancers include classical enhancers and facilitators to fully activate gene expression. Cell 186, 5826–5839.e18. 10.1016/j.cell.2023.11.030.

43. Kim, S., and Wysocka, J. (2023). Deciphering the multi-scale, quantitative cis-regulatory code. Molecular Cell 83, 373–392. 10.1016/j.molcel.2022.12.032.

44. Rayon, T., Menchero, S., Nieto, A., Xenopoulos, P., Crespo, M., Cockburn, K., Cañon, S., Sasaki, H., Hadjantonakis, A.-K., de la Pompa, J.L., et al. (2014). Notch and Hippo Converge on Cdx2 to Specify the Trophectoderm Lineage in the Mouse Blastocyst. Developmental Cell 30, 410–422. 10.1016/j.devcel.2014.06.019.

45. Rayon, T., Menchero, S., Rollán, I., Ors, I., Helness, A., Crespo, M., Nieto, A., Azuara, V., Rossant, J., and Manzanares, M. (2016). Distinct mechanisms regulate Cdx2 expression in the blastocyst and in trophoblast stem cells. Sci Rep 6, 27139. 10.1038/srep27139.

46. Watts, J.A., Zhang, C., Klein-Szanto, A.J., Kormish, J.D., Fu, J., Zhang, M.Q., and Zaret, K.S. (2011). Study of FoxA Pioneer Factor at Silent Genes Reveals Rfx-Repressed Enhancer at Cdx2 and a Potential Indicator of Esophageal Adenocarcinoma Development. PLoS Genet 7, e1002277. 10.1371/journal.pgen.1002277.

47. Benahmed, F., Gross, I., Gaunt, S.J., Beck, F., Jehan, F., Domon–Dell, C., Martin, E., Kedinger, M., Freund, J., and Duluc, I. (2008). Multiple Regulatory Regions Control the Complex Expression Pattern of the Mouse Cdx2 Homeobox Gene. Gastroenterology 135, 1238–1247.e3. 10.1053/j.gastro.2008.06.045.

48. Friman, E.T., Flyamer, I.M., Marenduzzo, D., Boyle, S., and Bickmore, W.A. (2023). Ultra-long-range interactions between active regulatory elements. Genome Res. 33, 1269–1283. 10.1101/gr.277567.122.

49. Montavon, T., and Duboule, D. (2013). Chromatin organization and global regulation of Hox gene clusters. Phil. Trans. R. Soc. B 368, 20120367. 10.1098/rstb.2012.0367.

50. Wang, W.C.H., and Shashikant, C.S. (2007). Evidence for positive and negative regulation of the mouseCdx2 gene. J. Exp. Zool. 308B, 308–321. 10.1002/jez.b.21154.

51. Strumpf, D., Mao, C.-A., Yamanaka, Y., Ralston, A., Chawengsaksophak, K., Beck, F., and Rossant, J. (2005). Cdx2 is required for correct cell fate specification and differentiation of trophectoderm in the mouse blastocyst. Development 132, 2093– 2102. 10.1242/dev.01801.

52. Metzis, V., Steinhauser, S., Pakanavicius, E., Gouti, M., Stamataki, D., Ivanovitch, K., Watson, T., Rayon, T., Mousavy Gharavy, S.N., Lovell-Badge, R., et al. (2018). Nervous System Regionalization Entails Axial Allocation before Neural Differentiation. Cell 175, 1105–1118.e17. 10.1016/j.cell.2018.09.040.

53. Argelaguet, R., Lohoff, T., Li, J.G., Nakhuda, A., Drage, D., Krueger, F., Velten, L., Clark, S.J., and Reik, W. (2022). Decoding gene regulation in the mouse embryo using single-cell multi-omics (Developmental Biology) 10.1101/2022.06.15.496239.

54. Blassberg, R., Patel, H., Watson, T., Gouti, M., Metzis, V., Delás, M.J., and Briscoe, J. (2022). Sox2 levels regulate the chromatin occupancy of WNT mediators in epiblast progenitors responsible for vertebrate body formation. Nat Cell Biol 24, 633– 644. 10.1038/s41556-022-00910-2.

55. Delás, M.J., Kalaitzis, C.M., Fawzi, T., Demuth, M., Zhang, I., Stuart, H.T., Costantini, E., Ivanovitch, K., Tanaka, E.M., and Briscoe, J. (2023). Developmental cell fate choice in neural tube progenitors employs two distinct cis-regulatory strategies. Developmental Cell 58, 3–17.e8. 10.1016/j.devcel.2022.11.016.

56. Coutaud, B., and Pilon, N. (2013). Characterization of a novel transgenic mouse line expressing Cre recombinase under the control of the *Cdx2* neural specific enhancer: A Cre-driver Line for the Caudal Neuroectoderm. genesis 51, 777–784. 10.1002/dvg.22421.

57. Tsakiridis, A., Huang, Y., Blin, G., Skylaki, S., Wymeersch, F., Osorno, R., Economou, C., Karagianni, E., Zhao, S., Lowell, S., et al. (2014). Distinct Wnt-driven primitive streak-like populations reflect *in vivo* lineage precursors. Development 141, 1209–1221. 10.1242/dev.101014.

58. Evans, R., O’Neill, M., Pritzel, A., Antropova, N., Senior, A., Green, T., Žídek, A., Bates, R., Blackwell, S., Yim, J., et al. (2021). Protein complex prediction with AlphaFold-Multimer (Bioinformatics) 10.1101/2021.10.04.463034.

59. Erceg, J., Pakozdi, T., Marco-Ferreres, R., Ghavi-Helm, Y., Girardot, C., Bracken, A.P., and Furlong, E.E.M. (2017). Dual functionality of *cis*-regulatory elements as developmental enhancers and Polycomb response elements. Genes Dev. 31, 590– 602. 10.1101/gad.292870.116.

60. Gisselbrecht, S.S., Palagi, A., Kurland, J.V., Rogers, J.M., Ozadam, H., Zhan, Y., Dekker, J., and Bulyk, M.L. (2020). Transcriptional Silencers in Drosophila Serve a Dual Role as Transcriptional Enhancers in Alternate Cellular Contexts. Molecular Cell 77, 324–337.e8. 10.1016/j.molcel.2019.10.004.

61. Halfon, M.S. (2020). Silencers, Enhancers, and the Multifunctional Regulatory Genome. Trends in Genetics 36, 149–151. 10.1016/j.tig.2019.12.005.

62. Ahituv, N., Zhu, Y., Visel, A., Holt, A., Afzal, V., Pennacchio, L.A., and Rubin, E.M. (2007). Deletion of Ultraconserved Elements Yields Viable Mice. PLoS Biol 5, e234. 10.1371/journal.pbio.0050234.

63. Dickel, D.E., Ypsilanti, A.R., Pla, R., Zhu, Y., Barozzi, I., Mannion, B.J., Khin, Y.S., Fukuda-Yuzawa, Y., Plajzer-Frick, I., Pickle, C.S., et al. (2018). Ultraconserved Enhancers Are Required for Normal Development. Cell 172, 491–499.e15. 10.1016/j.cell.2017.12.017.

64. Duarte, P., Brattig Correia, R., Nóvoa, A., and Mallo, M. (2023). Regulatory changes associated with the head to trunk developmental transition. BMC Biol 21, 170. 10.1186/s12915-023-01675-2.

65. Exelby, K., Herrera-Delgado, E., Perez, L.G., Perez-Carrasco, R., Sagner, A., Metzis, V., Sollich, P., and Briscoe, J. (2021). Precision of tissue patterning is controlled by dynamical properties of gene regulatory networks. Development 148, dev197566. 10.1242/dev.197566.

66. Garnett, A.T., Square, T.A., and Medeiros, D.M. (2012). BMP, Wnt and FGF signals are integrated through evolutionarily conserved enhancers to achieve robust expression of Pax3 and Zic genes at the zebrafish neural plate border. Development 139, 4220–4231. 10.1242/dev.081497.

67. Mukherjee, S., Luedeke, D.M., McCoy, L., Iwafuchi, M., and Zorn, A.M. (2022). SOX transcription factors direct TCF-independent WNT/β-catenin responsive transcription to govern cell fate in human pluripotent stem cells. Cell Reports 40, 111247. 10.1016/j.celrep.2022.111247.

68. Doni Jayavelu, N., Jajodia, A., Mishra, A., and Hawkins, R.D. (2020). Candidate silencer elements for the human and mouse genomes. Nat Commun 11, 1061. 10.1038/s41467-020-14853-5.

69. Jiang, J., Cai, H., Zhou, Q., and Levine, M. (1993). Conversion of a dorsal-dependent silencer into an enhancer: evidence for dorsal corepressors. The EMBO Journal 12, 3201–3209. 10.1002/j.1460-2075.1993.tb05989.x.

70. Kirov, N., Zhelnin, L., Shah, J., and Rushlow, C. (1993). Conversion of a silencer into an enhancer: evidence for a co-repressor in dorsal-mediated repression in Drosophila. The EMBO Journal 12, 3193–3199. 10.1002/j.1460-2075.1993.tb05988.x.

71. Weintraub, S.J., Prater, C.A., and Dean, D.C. (1992). Retinoblastoma protein switches the E2F site from positive to negative element. Nature 358, 259–261. 10.1038/358259a0.

72. Bessis, A., Champtiaux, N., Chatelin, L., and Changeux, J.-P. (1997). The neuron-restrictive silencer element: A dual enhancer/silencer crucial for patterned expression of a nicotinic receptor gene in the brain. Proc. Natl. Acad. Sci. U.S.A. 94, 5906–5911. 10.1073/pnas.94.11.5906.

73. Kallunki, P., Edelman, G.M., and Jones, F.S. (1998). The neural restrictive silencer element can act as both a repressor and enhancer of L1 cell adhesion molecule gene expression during postnatal development. Proc. Natl. Acad. Sci. U.S.A. 95, 3233–3238. 10.1073/pnas.95.6.3233.

74. Kehayova, P., Monahan, K., Chen, W., and Maniatis, T. (2011). Regulatory elements required for the activation and repression of the protocadherin-α gene cluster. Proc. Natl. Acad. Sci. U.S.A. 108, 17195–17200. 10.1073/pnas.1114357108.

75. Koike, S., Schaeffer, L., and Changeux, J.P. (1995). Identification of a DNA element determining synaptic expression of the mouse acetylcholine receptor delta-subunit gene. Proc. Natl. Acad. Sci. U.S.A. 92, 10624–10628. 10.1073/pnas.92.23.10624.

76. Cunningham, T.J., Kumar, S., Yamaguchi, T.P., and Duester, G. (2015). *Wnt8a* and *Wnt3a* cooperate in the axial stem cell niche to promote mammalian body axis extension. Developmental Dynamics 244, 797–807. 10.1002/dvdy.24275.

77. Ribes, V., Le Roux, I., Rhinn, M., Schuhbaur, B., and Dollé, P. (2009). Early mouse caudal development relies on crosstalk between retinoic acid,Shh and Fgf signalling pathways. Development 136, 665–676. 10.1242/dev.016204.

78. Savory, J.G.A., Edey, C., Hess, B., Mears, A.J., and Lohnes, D. (2014). Identification of novel retinoic acid target genes. Developmental Biology 395, 199–208. 10.1016/j.ydbio.2014.09.013.

79. Chatagnon, A., Veber, P., Morin, V., Bedo, J., Triqueneaux, G., Sémon, M., Laudet, V., d’Alché-Buc, F., and Benoit, G. (2015). RAR/RXR binding dynamics distinguish pluripotency from differentiation associated cis-regulatory elements. Nucleic Acids Research 43, 4833–4854. 10.1093/nar/gkv370.

80. Dupé, V., Davenne, M., Brocard, J., Dollé, P., Mark, M., Dierich, A., Chambon, P., and Rijli, F.M. (1997). In vivo functional analysis of the Hoxa *-1* 3′ retinoic acid response element (3′ RARE). Development 124, 399–410. 10.1242/dev.124.2.399.

81. Houle, M., Prinos, P., Iulianella, A., Bouchard, N., and Lohnes, D. (2000). Retinoic Acid Regulation of Cdx1: an Indirect Mechanism for Retinoids and Vertebral Specification. Molecular and Cellular Biology 20, 6579–6586. 10.1128/MCB.20.17.6579-6586.2000.

82. Houle, M., Sylvestre, J.-R., and Lohnes, D. (2003). Retinoic acid regulates a subset of Cdx1 function in vivo. Development 130, 6555–6567. 10.1242/dev.00889.

83. Kumar, S., and Duester, G. (2014). Retinoic acid controls body axis extension by directly repressing *Fgf8* transcription. Development 141, 2972–2977. 10.1242/dev.112367.

84. Berenguer, M., Meyer, K.F., Yin, J., and Duester, G. (2020). Discovery of genes required for body axis and limb formation by global identification of retinoic acid– regulated epigenetic marks. PLoS Biol 18, e3000719. 10.1371/journal.pbio.3000719.

85. Ghyselinck, N.B., and Duester, G. (2019). Retinoic acid signaling pathways. Development 146, dev167502. 10.1242/dev.167502.

86. Ruberte, E., Dolle, P., Krust, A., Zelent, A., Morriss-Kay, G., and Chambon, P. (1990). Specific spatial and temporal distribution of retinoic acid receptor gamma transcripts during mouse embryogenesis. Development 108, 213–222. 10.1242/dev.108.2.213.

87. Ang, H.L., and Duester, G. (1997). Initiation of retinoid signaling in primitive streak mouse embryos: Spatiotemporal expression patterns of receptors and metabolic enzymes for ligand synthesis. Dev. Dyn. 208, 536–543. 10.1002/(SICI)1097-0177(199704)208:4<536::AID-AJA9>3.0.CO;2-J.

88. Dollé, P., Ruberte, E., Leroy, P., Morriss-Kay, G., and Chambon, P. (1990). Retinoic acid receptors and cellular retinoid binding proteins: I. A systematic study of their differential pattern of transcription during mouse organogenesis. Development 110, 1133–1151. 10.1242/dev.110.4.1133.

89. Gillespie, R.F., and Gudas, L.J. (2007). Retinoic Acid Receptor Isotype Specificity in F9 Teratocarcinoma Stem Cells Results from the Differential Recruitment of Coregulators to Retinoic Acid Response Elements. Journal of Biological Chemistry 282, 33421–33434. 10.1074/jbc.M704845200.

90. Patel, N.S., Rhinn, M., Semprich, C.I., Halley, P.A., Dollé, P., Bickmore, W.A., and Storey, K.G. (2013). FGF Signalling Regulates Chromatin Organisation during Neural Differentiation via Mechanisms that Can Be Uncoupled from Transcription. PLoS Genet 9, e1003614. 10.1371/journal.pgen.1003614.

91. Semprich, C.I., Davidson, L., Amorim Torres, A., Patel, H., Briscoe, J., Metzis, V., and Storey, K.G. (2022). ERK1/2 signalling dynamics promote neural differentiation by regulating chromatin accessibility and the polycomb repressive complex. PLoS Biol 20, e3000221. 10.1371/journal.pbio.3000221.

92. Hu, Y., Kazenwadel, J., and James, R. (1993). Isolation and characterization of the murine homeobox gene Cdx-1. Regulation of expression in intestinal epithelial cells. Journal of Biological Chemistry 268, 27214–27225. 10.1016/S0021-9258(19)74240-9.

93. Kreimer, A., Ashuach, T., Inoue, F., Khodaverdian, A., Deng, C., Yosef, N., and Ahituv, N. (2022). Massively parallel reporter perturbation assays uncover temporal regulatory architecture during neural differentiation. Nat Commun 13, 1504. 10.1038/s41467-022-28659-0.

94. Sawada, S. (1994). A lineage-specific transcriptional silencer regulates CD4 gene expression during T lymphocyte development. Cell 77, 917–929. 10.1016/0092-8674(94)90140-6.

95. Siu, G., Wurster, A.L., Duncan, D.D., Soliman, T.M., and Hedrick, S.M. (1994). A transcriptional silencer controls the developmental expression of the CD4 gene. The EMBO Journal 13, 3570–3579. 10.1002/j.1460-2075.1994.tb06664.x.

96. Clark, E., and Peel, A.D. (2018). Evidence for the temporal regulation of insect segmentation by a conserved sequence of transcription factors. Development, dev.155580. 10.1242/dev.155580.

97. Copf, T., Schröder, R., and Averof, M. (2004). Ancestral role of *caudal* genes in axis elongation and segmentation. Proc. Natl. Acad. Sci. U.S.A. 101, 17711–17715. 10.1073/pnas.0407327102.

98. Faas, L., and Isaacs, H.V. (2009). Overlapping functions of *Cdx1*, *Cdx2*, and Cdx4 in the development of the amphibian Xenopus tropicalis. Developmental Dynamics 238, 835–852. 10.1002/dvdy.21901.

99. Martin, B.L., and Kimelman, D. (2009). Wnt Signaling and the Evolution of Embryonic Posterior Development. Current Biology 19, R215–R219. 10.1016/j.cub.2009.01.052.

100. Morales, A.V., De La Rosa, E.J., and De Pablo, F. (1996). Expression of the Cdx-B homeobox gene in chick embryo suggests its participation in rostrocaudal axial patterning. Dev. Dyn. 206, 343–353. 10.1002/(SICI)1097-0177(199608)206:4<343::AID-AJA1>3.0.CO;2-I.

101. Serafimidis, I., Rakatzi, I., Episkopou, V., Gouti, M., and Gavalas, A. (2008). Novel Effectors of Directed and Ngn3-Mediated Differentiation of Mouse Embryonic Stem Cells into Endocrine Pancreas Progenitors. Stem Cells 26, 3–16. 10.1634/stemcells.2007-0194.

102. Truett, G.E., Heeger, P., Mynatt, R.L., Truett, A.A., Walker, J.A., and Warman, M.L. (2000). Preparation of PCR-Quality Mouse Genomic DNA with Hot Sodium Hydroxide and Tris (HotSHOT). BioTechniques 29, 52–54. 10.2144/00291bm09.

103. Robinson, J.T. (2011). Integrative genomics viewer. correspondence 29.

104. Ewels, Philip A, Peltzer, Alexander, Fillinger, Sven, Patel, Harshil, Alneberg, Johannes, Wilm, Andreas, Garcia, Maxime Ulysse, Di Tommaos, Paolo, and Nahsen, Sven (2020). The nf-core framework for community-curatedbioinformatics pipelines. Nat Biotechnol 38, 271–278. 10.1038/s41587-020-0435-1.

105. Castro-Mondragon, J.A., Riudavets-Puig, R., Rauluseviciute, I., Berhanu Lemma, R., Turchi, L., Blanc-Mathieu, R., Lucas, J., Boddie, P., Khan, A., Manosalva Pérez, N., et al. (2022). JASPAR 2022: the 9th release of the open-access database of transcription factor binding profiles. Nucleic Acids Research 50, D165–D173. 10.1093/nar/gkab1113.

106. Tan, G., and Lenhard, B. (2016). TFBSTools: an R/bioconductor package for transcription factor binding site analysis. Bioinformatics 32, 1555–1556. 10.1093/bioinformatics/btw024.

107. Mirdita, M., Schütze, K., Moriwaki, Y., Heo, L., Ovchinnikov, S., and Steinegger, M. (2022). ColabFold: making protein folding accessible to all. Nat Methods 19, 679– 682. 10.1038/s41592-022-01488-1.

108. Paysan-Lafosse, T., Blum, M., Chuguransky, S., Grego, T., Pinto, B.L., Salazar, G.A., Bileschi, M.L., Bork, P., Bridge, A., Colwell, L., et al. (2023). InterPro in 2022. Nucleic Acids Research 51, D418–D427. 10.1093/nar/gkac993.

109. Jumper, J., Evans, R., Pritzel, A., Green, T., Figurnov, M., Ronneberger, O., Tunyasuvunakool, K., Bates, R., Žídek, A., Potapenko, A., et al. (2021). Highly accurate protein structure prediction with AlphaFold. Nature 596, 583–589. 10.1038/s41586-021-03819-2.

